# Structured demographic buffering: A framework to explore the environment drivers and demographic mechanisms underlying demographic buffering

**DOI:** 10.1101/2023.07.20.549848

**Authors:** Samuel J L Gascoigne, Maja Kajin, Shripad Tuljapurkar, Gabriel Silva Santos, Aldo Compagnoni, Ulrich K Steiner, Anna C Vinton, Harman Jaggi, Irem Sepil, Roberto Salguero-Gómez

**Affiliations:** Department of Biology, South Parks Road, University of Oxford, Oxford, United Kingdom; Department of Biology, Biotechnical Faculty, University of Ljubljana, Večna pot 111, 1000 Ljubljana, Slovenia; Biology Department, Stanford University, Stanford, CA, USA; National Institute of the Atlantic Forest (INMA), Santa Teresa, Espírito Santo, Brazil; Institute of Biology, Martin Luther University Halle-Wittenburg, Halle (Saale), Germany; German Centre for Integrative Biodiversity Research (iDiv) Halle-Jena-Leipzig, Leipzig, Germany; Institute of Biology, Freie Universität Berlin, Berlin, Germany; National Laboratory for Grassland & Agro-ecosystems, Lanzhou University, China

**Keywords:** environmental stochasticity, integral projection models (IPMs), life history strategies, stochastic demography

## Abstract

Environmental stochasticity is a key determinant of population viability. Decades of work exploring how environmental stochasticity influences population dynamics have highlighted the ability of some natural populations to limit the negative effects of environmental stochasticity, one of these strategies being demographic buffering. Whilst various methods exist to quantify demographic buffering, we still do not know which environment factors and demographic characteristics are most responsible for the demographic buffering observed in natural populations. Here, we introduce a framework to quantify the relative effects of three key drivers of demographic buffering: environment components (*e*.*g*., temporal autocorrelation and variance), population structure, and demographic rates (*e*.*g*., progression and fertility). Using Integral Projection Models, we explore how these drivers impact the demographic buffering abilities of three plant species with different life histories and demonstrate how our approach successfully characterises a population’s capacity to demographically buffer against environmental stochasticity in a changing world.

## INTRODUCTION

Understanding how populations minimise the negative effects of environmental stochasticity is central to ecology and evolution (Sutherland *et al*. 2013). A key prediction of life history theory is that increases in the temporal variance of demographic rates (*e.g*., rates of progression, stasis, retrogression and fertility) lead to reductions in a population’s stochastic growth rate (*λ*_*s*_) (Tuljapurkar 1982, 1989). In extreme cases, this demographic rate variance can lead to local extinction (May 1973; Saether *et al*. 1998; Lennartsson & Oostermeijer 2001; Bull *et al*. 2007; Melbourne & Hastings 2008). Critically, environmental stochasticity, a key driver of demographic rate variance (Jongejans *et al*. 2010), is projected to increase due to climate change (Urban 2015; Bathiany *et al*. 2018; Di Cecco & Gouhier 2018; Masson-Delmotte *et al*. 2021). Therefore, understanding the environment drivers and demographic mechanisms influencing the relationship between environmental stochasticity and population dynamics is both important and timely.

Three key considerations are needed to relate demographic rate variance to population dynamics. First, there are limits to the amount of variance that demographic rate can exhibit without driving a population to local extinction (Arthreya & Karlin 1971; May 1973). Second, the negative effects of demographic rate variance on population growth are exacerbated when the environment drivers impact the demographic rate(s) of highest importance (*i.e*., sensitivity) to *λ*_*s*_. However, the negative effect of demographic rate variance on *λ*_*s*_ can be reduced (or increased) when demographic rates covary negatively (or positively) (Tuljapurkar 1982, 1989), as demographic rates can compensate (amplify) for one another within a timestep. For example, demographic compensation may occur if instances of low adult survival happen concurrently with high adult reproduction, or *vice versa* (Sheth & Angert 2018). Third, environment-vital rate reaction norms can moderate the relationship between demographic rate variance and *λ*_*s*_ (King & Hadfield 2019; Bruijning *et al*. 2020). Following Jensen’s inequality (1906), convex (U-shaped) environment-demographic rate reaction norms result in a positive effect of demographic rate variance on *λ*_*s*_, whereas concave (∩-shaped) reaction norms lead to a negative effect (Drake 2005; Koons *et al*. 2009). These three key considerations regarding the impact of stochastic environments on population dynamics have produced key predictions in life history theory (Tuljapurkar *et al*. 2009; Sæther *et al*. 2013), conservation biology (Foley 1994; Higgins *et al*. 2000), and agriculture science (Lande *et al*. 1997; Mack 2000). However, these three considerations alone do not allow us to quantify a population’s ability to accommodate demographic rate variance; demographic buffering does.

Quantifying demographic buffering in natural populations has been a dynamic area of study in recent decades. The field has moved from regression-based approaches, where the deterministic elasticities (or sensitivities) of demographic rates with respect to *λ* are regressed against the coefficient of variation (or variance) of demographic rates (Pfister 1998; Morris & Doak 2004; further examples in Hilde *et al*. 2020), to a derivative-based approach that uses the summation of stochastic elasticities of variance, 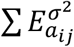, as a measure of demographic buffering (Santos *et al*. 2023; Wang *et al*. 2023). Despite important insights (*e.g*., McDonald *et al*. 2017), the regression-based approaches have important limitations, such as being confounded by the life cycle’s complexity, the lack of standardized methods (Hilde *et al*. 2020), and difficulty in clear-cut interpretations (see Santos *et al*. 2023 for further details).

Using the summation of stochastic elasticities of variance, one can explore the environment drivers and demographic mechanisms behind demographic buffering. This insight is possible because 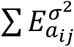 quantifies the proportional contribution of demographic rate variance to *λ*_*s*_ (Tuljapurkar *et al*. 2003; Haridas & Tuljapurkar 2005) and, consequently, directly quantifies degree of demographic buffering. Whilst researchers have previously used 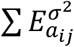 to quantify demographic buffering (Morris *et al*. 2008; Dalgleish *et al*. 2010), we still do not know how different environment components (*i.e*., temporal autocorrelation and variance), population structure (*i.e*., distribution of individuals in a population according to states, such as age, stage and/or size), and different demographic rates (*i.e*., state-specific transition probabilities or reproductive contributions between time *t* and *t* + 1) impact 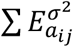.

Here, we test the effects of the environment components, population structure and demographic rates on the ability of natural populations to remain demographically buffered. We use environment-explicit stochastic integral projection models (IPMs) (Easterling *et al*. 2000; Ellner *et al*. 2016) for three perennial plant species from the PADRINO database (Levin *et al*. 2022) to test two hypotheses. We expect that: (H1) environment autocorrelation and variance will have negative effects on 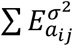. Specifically, as environments become more variable and positively autocorrelated, populations will become less buffered as predicted by Tuljapurkar’s (1982, 1989) small-noise approximation. (H2) Environment autocorrelation and variance influence 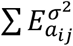 via different demographic mechanisms. Specifically, we expect that: (H2a) environment autocorrelation influences 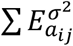 via its impact on population structure. We base this prediction on the fact that the impact of environment autocorrelation on population dynamics can be quantified by the degree to which the sequence of environments shifts the population from its long-term mean stable state structure (Tuljapurkar & Haridas 2006). Briefly, the rationale behind this expectation can be simplified by acknowledging that the commutative property of multiplication that applies to unstructured systems (*e.g*., 2 × 1 = 1 × 2) does not apply to structured systems (*e.g*., **A** × **B** ≠’ **B** × **A**, where **A** and **B** are matrices of size > 1 × 1). In turn, since the structure of the population is encoded into the population state distributions, we hypothesize that the impact of environment autocorrelation on 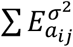 is strongly mediated by population structure. Similarly, we expect (H2b) environment variance to influence 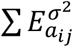 via the populations’ underlying demographic rates. This prediction also follows Tuljapurkar’s small-noise approximation (1982, 1989), where the impact of environment variance can be approximated by the summed product of the variance and sensitivities of individual demographic rates.

## METHODS

### Stochastic integral projection models

To explore the drivers of demographic buffering, we used integral projection models (IPMs). IPMs are discrete time population models (*i.e*., they project populations are projected across well-defined intervals of time from *t* to *t* + 1) that are structured with respect to a continuous variable (*e.g*., height, length, mass; Easterling *et al*. 2000; Ellner *et al*. 2016). To investigate the environment drivers and demographic mechanisms that impact degrees of demographic buffering in natural populations, we used environment explicit, parameter-stochastic IPMs for the *Berberis thunbergii* (Japanese barberry; Merow *et al*. 2017), *Calathea crotalifera* (rattlesnake plant; Westerband & Horvitz 2017) and *Heliconia tortuosa* (red twist Heliconia; Westerband & Horvitz 2017), extracted from the PADRINO IPM database (Levin et al. 2022). The chosen model structure allows us to individually influence regression parameters that underpin the IPM subkernels (*i.e*., the survival **P**- and fertility **F**-subkernels) based on the environment conditions to test our hypotheses.

We chose these three published IPMs to compare the roles of environment parameters and *λ*_*s*_ on 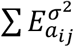 to gain some generality. The *B. thunbergii* IPM uses five environment parameters to build its kernels: mean temperature during warmest month, mean May precipitation, photosynthetically active radiation (PAR), soil nitrogen, and soil pH. The *C. crotalifera* and *H. tortuosa* IPMs use two environment parameters to define their kernels: canopy openness and photosynthetic rate. The kernel structure and parameters used in vital rate regressions for *B. thunbergii, C. crotalifera* and *H. tortuosa* are detailed in supplementary tables 1, 2 and 3, respectively. Furthermore, the models inhabit different domains of *λ*_*s*_. The models of *B. thunbergii* and *H. torutosa* have values of *λ*_*s*_ > 1 (*B. thunbergii*: *λ*_*s*_ = 1.378; *H. tortuosa*: *λ*_*s*_ = 1.367), implying long-term population growth, *C. crotalifera* has a *λ*_*s*_ < 1 (*λ*_*s*_ *=* 0.976), describing long-term population decline (Figure S1). Since *C. crotalifera* and *H. torutosa* have the same environment parameters and *B. thunbergii* and *H. tortuosa* have similar *λ*_*s*_values, by comparing demographic buffering across these species, we aim to examine possible impacts of environment parameters and *λ* on 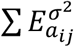 across the autocorrelation – proportional variance parameter space.

### Simulation methodology

To explore the roles of (H1) environment drivers as well as (H2a) population structure and (H2b) demographic rates on demographic buffering, we simulated IPMs across the environment autocorrelation – variance parameter space. In this simulation, all combinations of stochastic environment parameters, with autocorrelation ranging from -0.8 to 0.8 and proportional variance ranging from 0.9 (10% less variance in the environment than the IPM in PADRINO) to 1.1 (10% more variance in the environment that the IPM in PADRINO) were generated for all environment parameters. *B. thunbergii* had five environment parameters, whilst *C. crotalifera* and *H. tortuosa* had two environment parameters (Fig. 1a,b). We used these sequences of environment parameters to construct the time series of 1,000 IPM kernels from which we then estimated *λ*_*s*_ (eq. 1). Specifically, to calculate *λ*_*s*_: (1) a population of random structure was initialized, whereby the proportion of individuals of a given size class was generated from a uniform distribution ranging between the upper and lower limits of the IPMs (see Tables S1-3), (2) the population was then multiplied through the series of 1,000 parameter-stochastic IPM kernels, and (3) population sizes from timestep 200 to 1,000 were used to calculate *λ*_*s*_ following the equation:

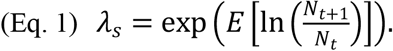

We omitted the first 200 projections from our calculation of *λ*_*s*_ to discard transient dynamics effects on short-term population size distributions (McDonald *et al*. 2016).

**Figure 1.**
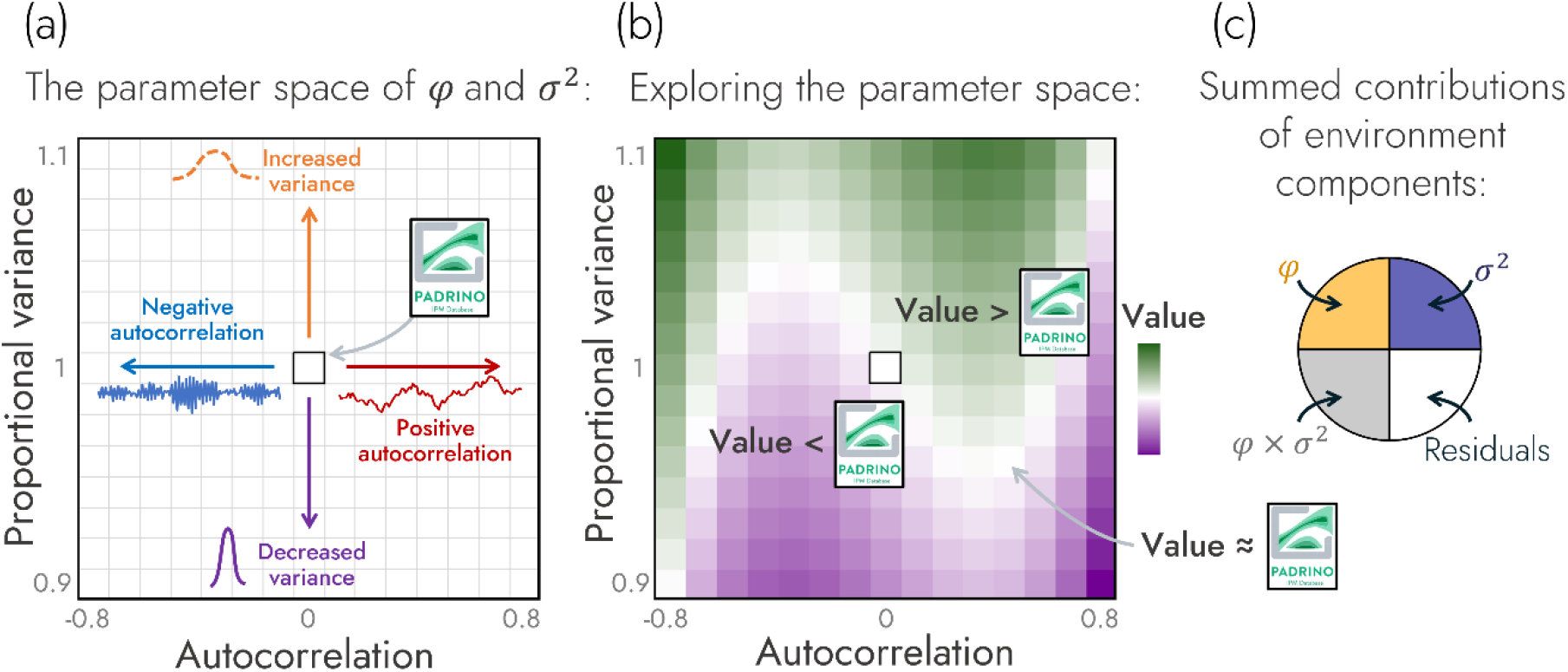
An overview of the simulation and analysis structure implemented to examine the impacts of climate drivers on natural populations. In our simulations, we explored how a population’s measure of demographic buffering changes over the parameter space of possible environment autocorrelation and variance values. (a) This space is visualized here across a 2D surface with environment autocorrelation on the x-axis and proportional variance on the y-axis. Environment variance is noted as proportional variance which is defined as the relative increase (>1) or decrease (<1) in the variance of a climate driver is made relative to the climate driver’s variance value stored in the PADRINO database. The middle of this landscape (*i.e*., autocorrelation = 0 and proportional variance = 1) represents the population model stored in the PADRINO database. (b) The impact of environment autocorrelation and variance on a response variable (*e.g*., degree of demographic buffering or a measure of population structure) is shown projected as a third dimension across this landscape. Across this projection, values lower than those reported in the original PADRINO IPM model are coloured purple, values close to the PADRINO model are coloured white, and values greater than the PADRINO model are coloured green. (c) The most parsimonious model that predicts the response variable as a function of environment autocorrelation and proportional variance was retained to calculate the summed linear and non-linear contribution of each predictor and the residuals towards the variance in the response variable.

### Generating environment time series

To explore the environment drivers of demographic buffering (H1), we manipulated the temporal autocorrelation and variance of environmental variables in our environmentally explicit stochastic IPMs. Whilst the effects of variance of demographic rates on population dynamics are commonly researched in population ecology (*e.g*., Jackson *et al*. 2022; Le Coeur *et al*. 2022), temporal autocorrelation is much less explored despite temporal autocorrelation having broad impacts on population dynamics (Petchey *et al*. 1997; Petchey 2000; Smallegange *et al*. 2014; Evers *et al*. 2023), life histories (Paniw *et al*. 2018; Vinton *et al*. 2023) and evolution (Wieczynski *et al*. 2018; Vinton *et al*. 2022). To fill this gap in knowledge, we used a first-order autoregressive function to generate the sequence of environment values used to build the series of IPM kernels. Here, *φ* represents the degree of autocorrelation across time steps whilst, *ϵ*_*t*+1_ represents white noise (*i.e*., random draws from a normal distribution, **ϵ**∼*N*(0,1)).

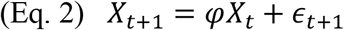

Subsequently, to coerce the autocorrelated series (**X**) to realistic values for the vital rate regressions that build the IPMs (shown in Tables S1-3), the final sequence of environment values was to a desired mean (*μ*) and variance (*σ*^2^) of the simulated environment:

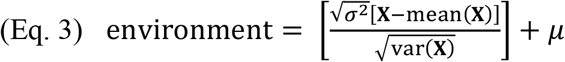

As our objective is not to evaluate the effect of shifts in mean environment values on demographic buffering but rather to examine the impacts of variance and autocorrelation, *μ* values were kept constant across simulations, whilst *σ*^2^ values varied across simulations.

Since the environment variables across the three species have different variances 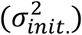, to standardize the increase/decrease in environment variance across parameters, we manipulated variances proportional to their variances coded in the PADRINO database 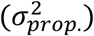 (Levin *et al*. 2022).

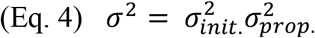

### Analysing the effects of environment autocorrelation and variance

To explore the effects of environmental components on each species’ ability to remain demographically buffered (H1,2), we constructed a suite of linear models using autocorrelation and proportional variance as predictors whilst also including an autocorrelation × proportional variance as an interaction term. Furthermore, since the impact of autocorrelation and proportional variance on demographic buffering may be nonlinear, we also constructed models using the quadratic and cubic forms of proportional variance and autocorrelation as predictors. To select the most appropriate model to describe the data, we used model comparison based on AIC (see supplementary materials p. 4 for the full analysis pipeline and Tables S4-12 for full AIC break down). After selecting the most parsimonious model, we calculated the proportion of variance in 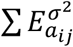 that can be explained by the summed contributions of autocorrelation, proportional variance, autocorrelation × proportional variance and residuals (Figure 1c).

### Perturbation analyses to quantify 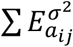

To quantify the degree of demographic buffering across our simulations (testing H1,2), we calculated the summation of stochastic elasticities of variance of demographic rates with respect to *λ*_*s*_. We estimated this variable, 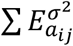, *numerically*. Whilst the **K**-kernel of an IPM is defined as a continuous map that projects a continuously structured population across time steps, in practice we discretise the kernel into a matrix notated as **A** (Easterling *et al*. 2000; Ellner *et al*. 2016). Since **A** is composed of individual matrix elements (*a*_*ij*_) and our stochastic environment generates a temporal sequence of **A** matrices, we can quantify the temporal variance of each *a*_*ij*_ element in matrix **A**. In turn, we numerically calculate 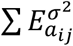 by perturbing the temporal variance of each matrix element (*a*_*ij*_) from our IPMs individually by 0.00001 proportionate (elasticity) to the unperturbed temporal variance of that matrix element. After perturbation of the matrix element, we calculated a perturbed stochastic population growth rate 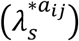. The summation of these weighted differences in *λ*_*s*_ and 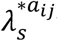 yields 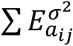.

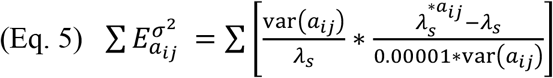

To calculate the impact of demographic rates on demographic buffering (H2b), we perturbed the subkernels that describe survival-dependent changes in size (**P**) and fertility (**F**) using the same method we used for the **K**-kernels. After calculating the subkernel-level elasticities of variance (Griffith 2017), we subtracted the subkernel summed elasticities of demographic rates to calculate their relative contributions: **P – F** contribution. Positive (negative) values of **P – F** contribution indicate relative variance in rates of survival-dependent changes in size are more (less) impactful on *λ*_*s*_ than relative variance in rates of fertility.

### Quantifying the impact of population structure on 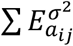

To analyse how population structure influences demographic buffering (H2a), we used two numerical approaches. Whilst methods exist to *analytically* measure the impact of population structure on asymptotic properties of population dynamics (Tuljapurkar & Lee 1997), currently there are no analytical approaches to quantify the degree to which multiple environment components influence 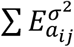 via population structure. In turn, we use two measures of population structure using a *regression-based approach* and an *estimate-based approach*. These approaches *numerically* link the impact of environment autocorrelation and variance on 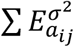 via population structure. Importantly, using these two approaches to investigate H2a allows us to cross-validate outputs (*i.e*., the hypothesized result of environment autocorrelation impacting 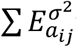 via shifts in population structure).

The *regression-based approach* involved examining deviances from stationary distributions. To do so, we regressed the scaled values – relative to the average size distribution – of the expected mean buffering value of a randomly selected individual in the population 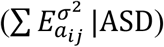 against scaled values of 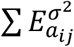. Deviances of 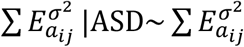 from a 1-to-1 line (*i.e*., the existence of residuals from this regression) indicates shifts in population structure may be influencing 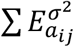. Subsequently, regressing these residuals against the environment components allows us to implicate an environment component – hypothesized to be environment autocorrelation [H2a] – as driving the impact of population structure on 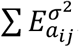. To perform this approach, we weighted 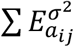 by the average size distribution (*i.e*., the average size distribution [ASD] of individuals in the population across the simulation) to calculate 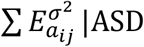. To determine the population’s average size distribution for a given environment, we iterated 1,000 randomly generated size distributions through the series of stochastic kernels and retained the mean of all size distributions across time steps 200 to 1,000 as an estimation of the average size distribution. Burning in the first 200 timesteps mitigates the impact of transients on the ASD. After calculating 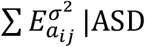, the emergent distribution was z-transformed (mean = 0, standard deviation = 1) and regressed against z-transformed values of 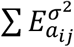 not informed by the average size distribution. Residuals from this regression represent a possible impact of population structure on 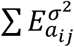. To further investigate the impact of environment autocorrelation and variance on 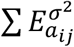 via said residuals, we modelled the residuals of the 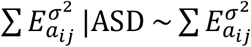 regression in response to environment autocorrelation and variance.

The *estimate-based approach* involved calculating the mean of the distribution of demographic buffering across a life history, termed *mean buffered size*. Calculating mean buffered size allows us to explore if the degree of buffering across a life history is shifted towards smaller or larger sizes across the environment autocorrelation – variance parameter space. To calculate this mean buffered size, we calculated the relative size (*i.e*., 0 = smallest possible size (α) and 1 = maximum possible size (ω)) that corresponds to the centre of the distribution of 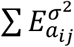 across the domain of sizes (Eq. 6). This calculation mirrors the method of calculating generation time as the mean age of reproductive individuals in the population (Ebert 1999, pg. 14).

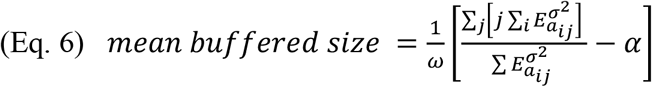

After calculating the mean buffered size for each species across the environment autocorrelation – variance parameter space, we regressed mean buffered size against the environment components to test our hypothesis that environment autocorrelation influences 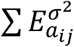 via shifts in population structure (H2a).

## RESULTS

### Testing H1: Environment variance is the primary driver of demographic buffering

Here we tested the hypothesis that environment autocorrelation and variance have negative effects on demographic buffering as quantified via 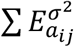 (H1). To do so, we ran simulations of the *Berberis thunbergii, Calathea crotalifera* and *Heliconia tortuosa* IPMs across the domain of autocorrelation and proportional variance values and calculated 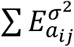. We found environment variance to be the primary driver of variance in 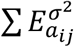 (Figure 2). The summed contributions of proportional variance accounted for 94% of the variance of 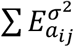 in *B. thunbergii* (R^2^ = 0.99, Table S4) (Figure 2a), 85% of the variance of 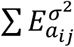 in *C. crotalifera* (R^2^ = 0.89, Table S5 (Figure 2b) and 83% of the variance of 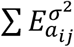 in *H. tortuosa* (R^2^ = 0.89, Table S6) (Figure 2c). Supporting our hypothesis, environment variance had a negative effect on 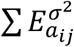 (see models for *B. thunbergii, C. crotalifera*, and *H. tortuosa* in Tables S4-6). However, we did not find evidence for a negative effect of environment autocorrelation on 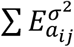. Instead, all species were best modelled when the quadratic and cubic forms of autocorrelation were used as predictors of 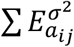 without the inclusion of a linear effect of autocorrelation. This finding indicates the impact of autocorrelation on 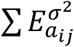 is non-linear across the environment autocorrelation and variance parameter space.

**Figure 2.**
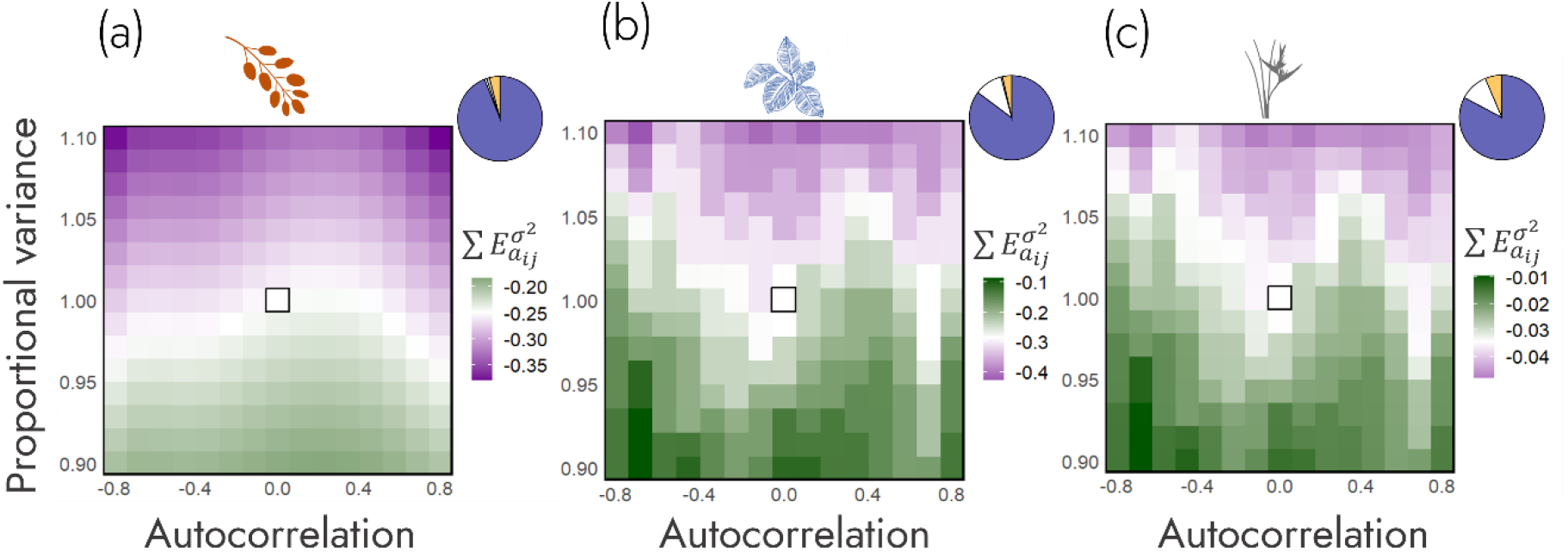
Environment variance (*σ*^2^) is the primary driver of demographic buffering. Across *Berberis thunbergii* (a), *Calathea crotalifera* (b) and *Heliconia tortuosa* (c), environment variance (blue in pie-chart) explains the majority of variance in 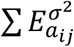. Populations of all three species become relatively less buffered (lower values of 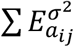, in purple) as proportional variance of environment components increase, whilst populations become relatively more buffered (higher values of 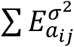, in green) as environment variance decreases. This strong impact of proportional variance of environment components is summarized in the pie charts detailing the proportion of variance in 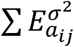 that can be explain by the environment components: environment autocorrelation in orange, environment variance in blue, environment autocorrelation × variance interaction in grey (so small here it is not visible), and unexplained residuals in white. Since the pie charts are predominantly blue across all three species, variance in environment components is the primary driver of 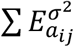 across the environment autocorrelation – variance parameter space.

### Testing H2a: Temporal autocorrelation influences demographic buffering via population structure

We used two approaches to test the hypothesis that temporal autocorrelation influences demographic buffering via shifts in population structure (H2a). First, we used a measure of demographic buffering that accounts for population structure 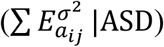 and regressed that against our normal measure of demographic buffering 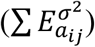. Second, we measured the shifts in the distribution of buffering across the life history in response to environment components.

In our first approach, we regressed scaled values of 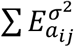 across all simulations against their respective 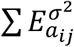 normalized by simulation specific stable size distribution 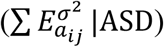. Since both values are scaled to mean = 0 with standard deviation = 1, any deviation of 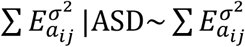 from the 1-to-1 regression line indicates temporal shifts in population structure may impact demographic buffering. Interestingly, we found heterogeneity in the degree to which 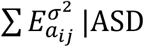 differed from 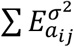 across species. Whilst *C. crotalifera* reported a 1-to-1 regression line between 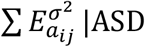 and 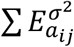 (R^2^ = 1, Figure 3d), *B. thunbergii* and *H. tortuosa* had residuals (*B. thunbergii*: R^2^ = 0.9977, Fig. 3a; *H. tortuosa*: R^2^ = 0.9995, Figure 3g). These residuals indicate that population structure may influence 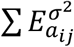, specifically in *B. thunbergii* and *H. tortuosa*.

**Figure 3.**
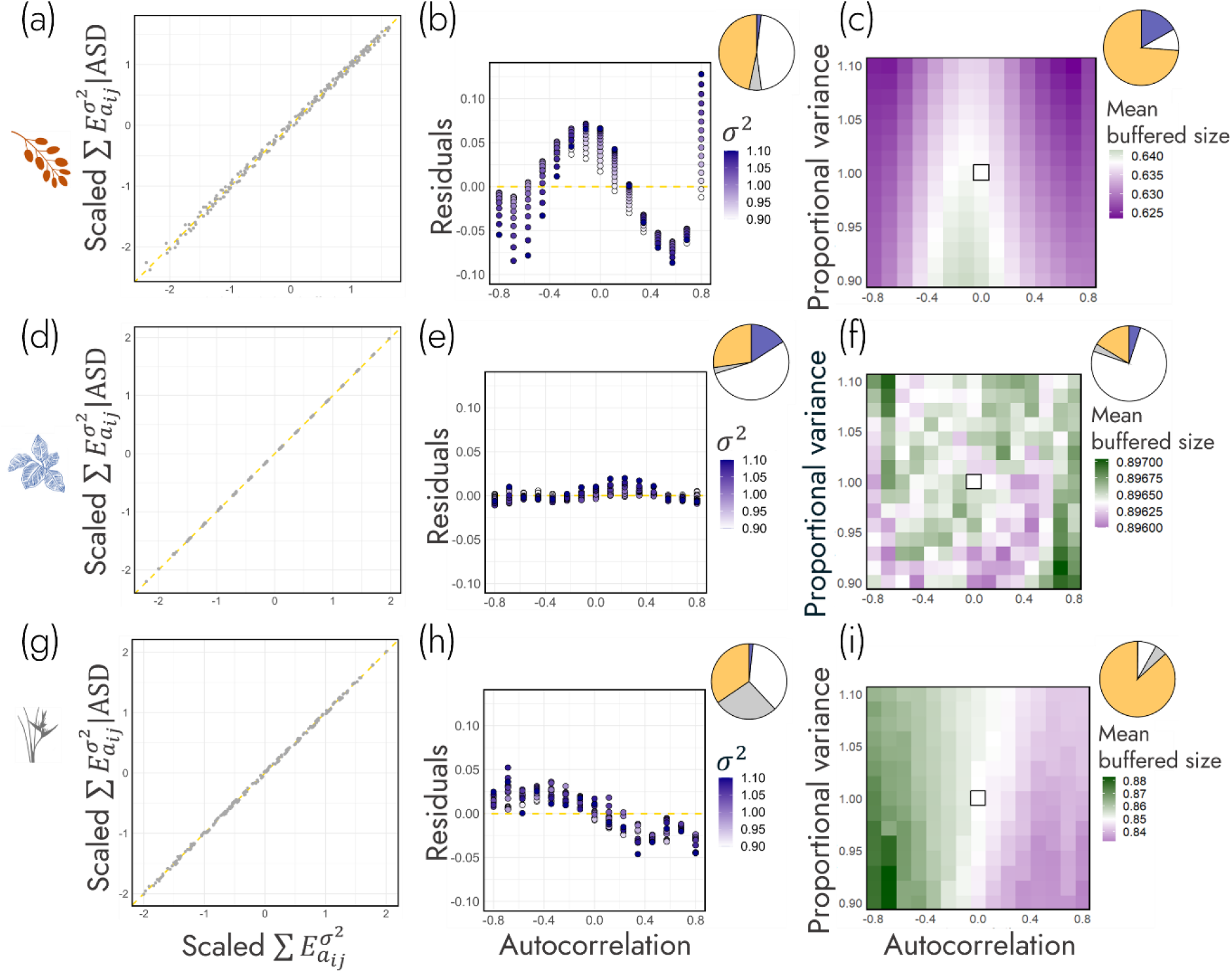
Environment autocorrelation can influence demographic buffering 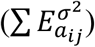 via its impact on population structure. In addition, the degree to which environmental autocorrelation impacts 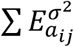 across *Berberis thunbergii* (a-c), *Calathea crotalifera* (d-f) and *Heliconia tortuosa* (g-i) is species-specific. The first column (a, d, g) shows the correlation between 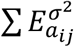 and demographic buffering weighted by the average stage distribution 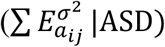. Residuals from these regressions show the potential impact of population structure on 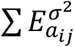. We then, in the second column (b, e, h), investigate these residuals as a function of the environment autocorrelation (x-axis) and environmental variance (*σ*^2^; purple). Lastly, in the third column (c, f, i), we quantify the impact of environment autocorrelation and variance on the mean buffered size of the population. The pie charts at the top right-hand corner of panels in (b, e, h), and (c, f, i) detail the proportion of variance in 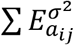 that is explained by environment autocorrelation (orange), environment variance (blue), environment autocorrelation × variance interaction (grey) and residuals (white). These pie charts show how environmental autocorrelation is the primary driver of shifts in 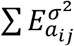 due to population.

To determine if environment autocorrelation is driving these residuals, we modelled the residuals of the 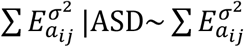 regression against environment autocorrelation and variance. Supporting our hypothesis (H2a), we found the residuals of the 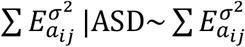 regression are mostly explained by environment autocorrelation (Figures 3b,e,h). In *B. thunbergii* and *H. tortuosa* (the species with the largest residuals from the 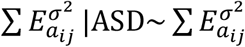 regression), environment autocorrelation accounted for 48% (R^2^ = 0.56, Figure 3b, Table S7) and 46% (R^2^ = 0.84, Figure 3h, Table S9) of the variance in residuals respectively; whilst environment variance only accounted for 2% of the variance in residuals in both species. Regarding *C. crotalifera*, the largest contributor to variance in residuals was unexplained residual variance (56%, R^2^ = 0.47, Figure 3e, Table S8), followed by environment autocorrelation (28%) and variance (16%).

In our second approach, we analysed the impact of environment autocorrelation and variance on the distribution of demographic buffering across a life cycle. In turn, we calculated the centre of the distribution of demographic buffering across a life history: mean buffered size. Echoing the findings from the first line of enquiry, mean buffered size was best explained by changes in environment autocorrelation – especially in *B. thunbergii* and *H. tortuosa*. Specifically, in *B. thunbergii*, 73% of the variance in mean buffered size was attributed to environment autocorrelation whilst 17% was attributed to environment variance (R^2^ = 0.91, Figure 3c, Table S10). Additionally, in *H. tortuosa*, 91% of the variance in mean buffered size was attributed to environment autocorrelation with only 0.1% being attributed to changes in environment variance (R^2^ = 0.97, Figure 3i, Table S12). And finally, just as in the first line of enquiry, 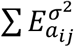 in *C. crotalifera* is less exposed to impacts of shifts in population structure as the distribution of mean buffered size across the environment autocorrelation – variance parameter space was mostly explained by residual variance (78%) rather than environment autocorrelation (17%) or environment variance (5%) (R^2^ = 0.26, Figure 3f, Table S11).

### Testing H2b: Demographic buffering is most sensitive to environment variance’s impact on rates of progression

To test the hypothesis that environment variance impacts demographic buffering through vital rates (H2b), we ran the same perturbation analysis used to calculate 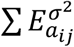 at the level of the sub-kernels: **P**-subkernel (survival-dependent changes in size) and the **F**-subkernel (fertility). By taking the difference of the subkernel elasticities of variance (*i.e*., **P – F** contribution), we investigated (1) the role of underlying rates on demographic buffering and (2) the environmental components that influence the **P – F** contribution across the environment autocorrelation – variance parameter space.

First, we determined if the **P – F** contribution is a sufficient predictor of 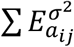. The **P – F** contribution was highly predictive of 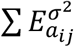 across all species (Figure 4a). *B. thunbergii* had a negative relationship between **P – F** contribution and 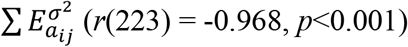, whilst *C. crotalifera* and *H. tortuosa* had positive relationships (*C. crotalifera*: *r*(223) = 0.999, *p*<0.001; *H. tortuosa*: *r*(223) = 0.983, *p*<0.001). These results indicate lower degrees of demographic buffering are associated with a greater impact of variance in rates of progression (*vs*. fertility) in *B. thunbergii*, but the opposite, a greater impact of variance in fertility (*vs*. progression) in *C. crotalifera* and *H. tortuosa*.

**Figure 4.**
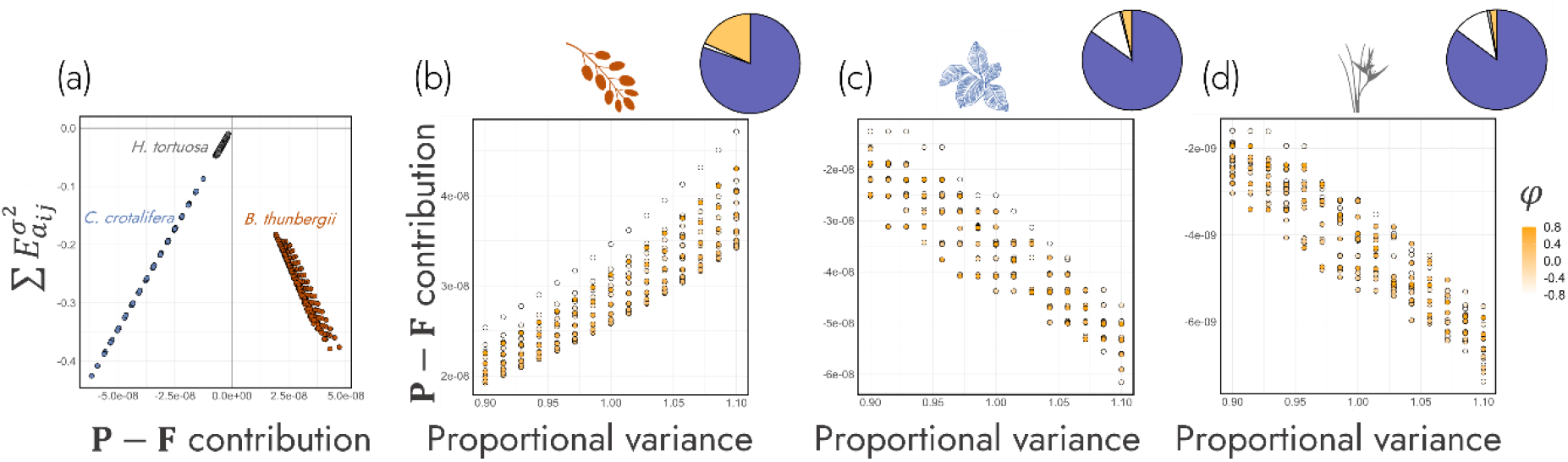
Environment variance (*σ*^2^) influences demographic buffering 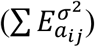 via the population’s underlying demographic rates. (a) The relative contribution of progression (growth conditional on survival: **P**) and fertility (recruitment of new individuals from reproductive ones the previous year: **F**) on 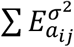 (*i.e*., **P**-**F** contribution). This approach was then applied to three plant species: (b) *Berberis thunbergii*, (c) *Calathea crotalifera*, and (d) *Heliconia tortuosa*). Dots are coloured by the degree of environment autocorrelation (yellow). The pie charts at the top right-hand corner of panels b-d detail the proportion of variance in 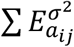 that is explained by environment autocorrelation (*φ*, orange), environment variance (blue), environment autocorrelation × variance interaction (grey) and residuals (white). These pie charts show how environment variance is the primary driver of shifts in the relative contributions of progression and fertility to 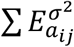.

To test if variance in **P – F** contribution is most explained by environment variance rather than autocorrelation (H2b), we regressed **P – F** contribution against the environment components. Across the three species, the **P – F** contribution was mostly explained by differences in degrees of environment variance rather than autocorrelation across the environment autocorrelation – variance parameter space (Figures 4b-d). Specifically, environment variance explained 80%, 85% and 86% of the variance of **P – F** contribution in *B. thunbergii* (R^2^ = 0.99, Figure 4b, Table S13), *C. crotalifera* (R^2^ = 0.89, Figure 4c, Table S14) and *H. tortuosa* (R^2^ = 0.89, Figure 4d, Table S15), respectively. However, of the remaining variance, environment autocorrelation explained 17%, 3% and 2% of the variance of **P – F** contribution, respectively.

## DISCUSSION

Environment drivers and demographic mechanisms are key to quantify and predict a population’s capacity for demographic buffering. Using three stochastic IPMs from the PADRINO database (Levin *et al*. 2022), we obtain partial support for the hypothesis that environment autocorrelation and variance negatively impact a population’s capacity to remain demographically buffered (H1). Interestingly, whilst environment variance negatively affects demographic buffering, there is a nonlinear effect of temporal autocorrelation on demographic buffering. Furthermore, even though environment autocorrelation and variance combine to make the environment time series, we show that their effects on demographic buffering are orthogonal dimensions of environmental stochasticity. Indeed, the effect of temporal autocorrelation on demographic buffering 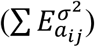 is mediated by population structure (H2a), whilst the effect of environment variance on 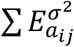 is mediated by underlying demographic rates (H2b). Specifically, the influence of environment variance on rates of progression *vs*. fertility is the greatest driver of differences in 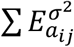 across variable environments in the three examined species. This finding builds on multiple lines of evidence showing how different life histories can persist in variable environments via the differential variance of progression *vs*. fertility rates (Gaillard *et al*. 1998; Pfister 1998).

Identifying the mechanisms that underpin the ability of natural populations to buffer against environmental stochasticity offers a powerful framework to explore a population’s vulnerability to climate change. Current climatic forecasts predict environmental stochasticity to increase with global climate change (Masson-Delmotte *et al*. 2021). For example, periods of extreme variation in temperature and precipitation are expected to increase in the tropics and sub-tropics which host the highest biodiversity (temperature: Bathiany *et al*. 2018; precipitation: Trenberth 2011). Furthermore, extreme weather events are expected to become more common, leading to increased autocorrelation (*e.g*., tropical cyclones: Knutson *et al*. 2010; fire frequency: Halofsky *et al*. 2020). However, not all environmental components affect populations the same way (Hoffmann & Bridle 2022; Vinton *et al*. 2022, 2023). The shape of demographic rates across a life history varies widely across the tree of life (Jones *et al*. 2014; Salguero-Gómez *et al*. 2017; Paniw *et al*. 2018; Healy *et al*. 2019; Varas-Enriquez *et al*. 2022). Therefore, predicting the susceptibility of populations to environmental stochasticity, without a regard to the mechanism, overlooks key heterogeneity in the demographic processes necessary for accurate predictions. Our framework provides a promising avenue to incorporate this heterogeneity for informed analyses of the role of environmental stochasticity in a population’s demographic buffering capacity.

Our results highlight an interesting, but often overlooked, role of population structure in demographic buffering. Whilst we find environment autocorrelation to primarily impact demographic buffering via shifts in population structure, there is also species-level heterogeneity in the strength and direction by which environment autocorrelation shifts population structure. Furthermore, our results indicate portions of the heterogeneity in 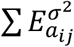 are explained by the interaction between environment autocorrelation and variance. One likely source of this heterogeneity is transient dynamics (*i.e*., short-term, progressively weakening realizations of non-asymptotic lambda values resulting from a population not being at its stable-stage distribution (Stott *et al*. 2011)). Whilst transient dynamics represent a suite of different stereotyped population dynamics (Capdevila *et al*. 2020), only *reactivity* (the degree to which a population not at its stable-stage distribution increases/decreases relative to that same population projected from its stable-stage distribution (Neubert & Caswell 1997)) has been linked to stochastic demography (McDonald *et al*. 2016). However, the link between reactivity, along with other transient dynamics, and demographic buffering remains unknown. Future work analysing which transient dynamics are increasing and decreasing levels of demographic buffering will finally integrate the analysis of transient dynamics with stochastic demography.

Historically, studies of life histories in stochastic environments have followed two branches: modelling and dimension reduction. Modelling life histories in stochastic environments, whereby analytic or numeric methods are used for demographic inference in individual populations, has progressively put to rest some key problems within life history theory (iteroparity: Orzack & Tuljapurkar 1989; Tuljapurkar *et al*. 2009; diapause: Tuljapurkar & Istock 1993; migration: Wiener & Tuljapurkar 1994; biennialism: Klinkhammer & de Jong 1983; Roerdink 1988, 1989; homeostasis: Orzack 1985; lability: Koons *et al*. 2009; Jongejans *et al*. 2010; Barraquand & Yoccoz 2013; summarized in Caswell (2001, pg. 440)). However, one of the limitations of a modelling approach is losing the realism captured within constraints, phylogenetic history or selection gradients that drive variance patterns in demographic rates.

From the empirical side, researchers have used dimension reduction techniques to unmask the patterns life histories exhibit in variable environments. Dimension reduction techniques, such as phylogenetically controlled principal component analyses (Revell 2012), are especially useful as a life history is not a value nor an object; a life history strategy is an abstract concept that researchers probe with life history traits – such as: longevity, age at maturity, average body size, *etc*. To capture the signal of an individual life history strategy through the dimensionality, reducing the multidimensionality of life history metrics to its most important axes of variance (*i.e*., principal components) has led to key discoveries (two-axes of life history variance: Salguero-Gómez *et al*. 2017; Healy *et al*. 2019). Furthermore, this approach has been used to model life histories in stochastic environments (Paniw *et al*. 2018; Romeijn & Smallegange 2022). However, this approach is limited to modelling only one component of a variable environment (*e.g*., environment autocorrelation *or* variance). This limitation is further emphasized by our results showing non-linearities between the effects of environmental components on 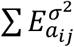, thereby illustrating that the impact of an environment component on demographic process is context dependent.

Using our framework, researchers can stitch the modelling and dimension reduction approaches together. Our framework can be applied to any environmentally explicit structured population models: from physiologically structured population models (de Roos 1997) to matrix population models (Caswell 2001) to integral projection models (Easterling *et al*. 2000; Ellner *et al*. 2016), to dynamic energy budget models (Nisbet *et al*. 2000; Smallegange *et al*. 2017). By using open-access data (COMPADRE: Salguero-Gómez *et al*. 2015; COMADRE: Salguero-Gómez *et al*. 2016; PADRINO: Levin *et al*. 2022; AmP: Marques *et al*. 2018), researchers can explore the combined impact of autocorrelation and variance on 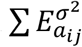 by interfacing the time series of a structured population models with stochastic matrices (as in Paniw *et al*. 2018). Once the landscape of 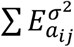 is mapped across environment autocorrelation and variance, the relative contributions of constraints, phylogeny and species-specific effects on 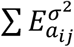 will be realized. This combined approach of modelling and dimension reduction offers generalization in a previously exception driven area of life history theory.

In conclusion, structure matters. Since Leslie (1945) and Lefkovitch (1965), demographers have explored how relatively simple structured population models can be used for biological inference. From transient dynamics (Hastings 2001; Ezard *et al*. 2010; Capdevila *et al*. 2020, 2022), to structured Lotka-Volterra models (de Roos *et al*. 1990; de Roos 2021) to stability analysis (Cushing *et al*. 2003), researchers have generated a rich body of theory and evidence for the impact of population structure on demographic inferences. However, the impact of environment structure, in the form of individual climate drivers (*e.g*., temporal autocorrelation and variance), and their corresponding demographic mechanisms that mediate their effects are uncoupled. We argue they should be stitched together. Our framework exploring demographic buffering across the environment autocorrelation – variance parameter space joins a recent push stitching the impacts of climate drivers (*e.g*., Vinton *et al*. 2022) with their respective demographic mechanisms (*e.g*., Le Coeur *et al*. 2022).

## ACKNOWLEDGMENTS

We thank Christina M. Hernández, for feedback on a previous version of this manuscript. M.K. was supported by a Marie Curie Fellowship (MSCA MaxPersist #101032484) hosted by R.S-G.; G.S.S. was supported by CNPq (#301343/2023-3); A.C was funded by the DFG (Deutsche Forschungsgemeinschaft #506492810). U.K.S was funded by the German Science Foundation (DFG Project #430170797). A.C.V. was supported by the National Science Foundation Postdoctoral Research Fellowship (#2010783) hosted by R.S-G. and I.S.; I.S. was supported by a Biotechnology and Biological Sciences Research Council (BBSRC) Fellowship (#BB/T008881/1), a Royal Society Dorothy Hodgkin Fellowship (#DHF\R1\211084), and a Wellcome Institutional Strategic Support Fund, University of Oxford (#BRR00060); R.S-G. was supported by a NERC Independent Research Fellowship (#NE/M018458/1).

## Notes

<u>Data accessibility statement:</u> All data and code supporting these results will be made open-access on Zenodo upon publication.

### Competing Interest Statement

The authors have declared no competing interest.

