## Supplementary online materials for "Structured demographic buffering: A framework to explore the environment drivers and demographic mechanisms underlying demographic buffering"

Samuel J L Gascoigne[1,*], Maja Kajin[1,2], Shripad Tuljapurkar[3], Gabriel Silva Santos[4], Aldo Compagnoni[5,6], Ulrich K Steiner[7], Anna C Vinton[1], Harman Jaggi[3], Irem Sepil[1] & Roberto Salguero-Gómez [1,8]

[1] Department of Biology, 11a Mansfield Rd, University of Oxford, Oxford, United Kingdom
[2] Department of Biology, Biotechnical Faculty, University of Ljubljana, Večna pot 111, 1000 Ljubljana, Slovenia
[3] Biology Department, Stanford University, Stanford, CA, USA
[4] National Institute of the Atlantic Forest (INMA), Santa Teresa, Espírito Santo, Brazil
[5] Institute of Biology, Martin Luther University Halle-Wittenburg, Halle (Saale), Germany
[6] German Centre for Integrative Biodiversity Research (iDiv) Halle-Jena-Leipzig, Leipzig, Germany
[7] Institute of Biology, Freie Universität Berlin, Berlin, Germany
[8] National Laboratory for Grassland & Agro-ecosystems, Lanzhou University, China

***** corresponding

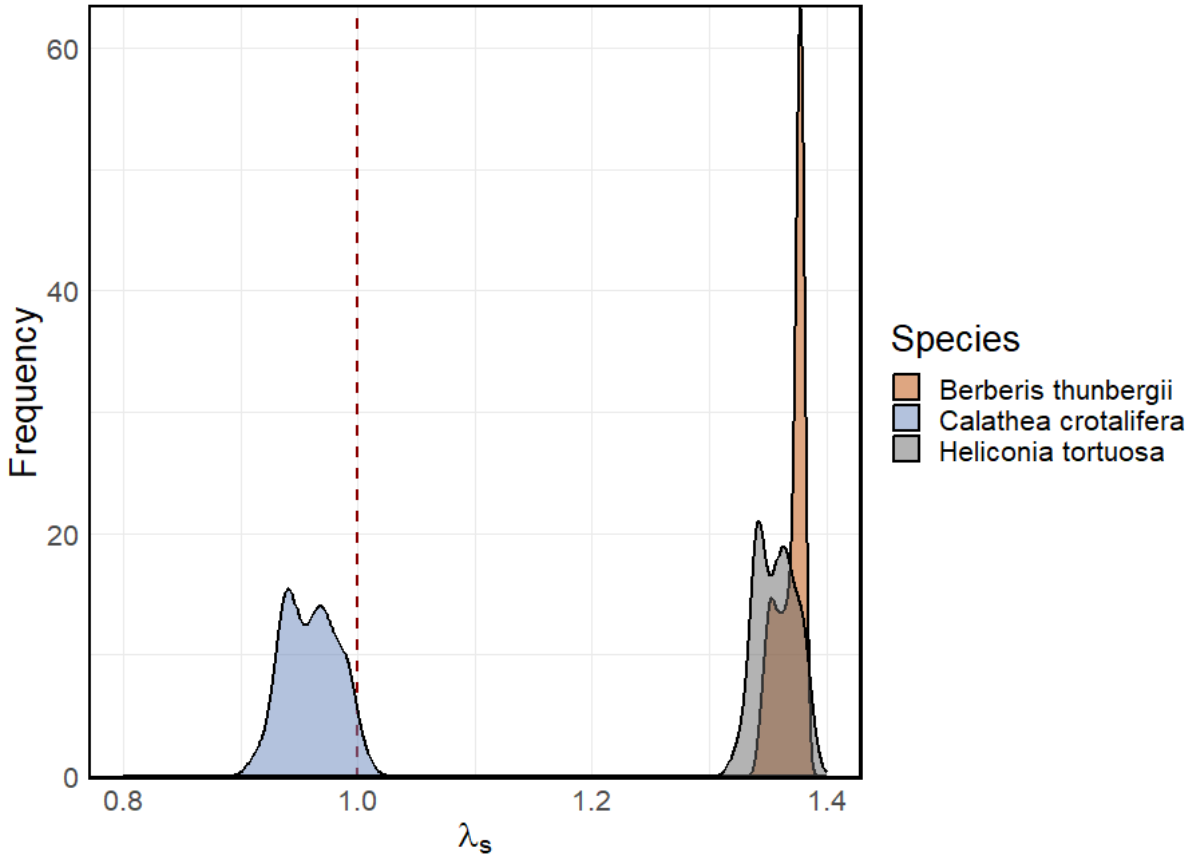

**Supplementary Figure 1**. Distribution of $\lambda_s$ from simulations across the environment autocorrelation – variance parameter space. The red dashed line represents a population that stays stable when projected across time steps (*i.e.*, $\lambda_s = 1$). Values distributed to the left of the red dashed line represent populations who will asymptotically decline in number across time steps (*i.e.*, $\lambda_s < 1$), whilst values distributed to the right of the red dashed line represent populations that will asymptotically increase across time steps (*i.e.*, $\lambda_s > 1$).

**Supplementary Table 1.** Formulas, regressions and parameters used to construct the IPMs for *Berberis thunbergii*.

| Construction | | Model | Parameter |
|---|---|---|---|
| Density-independent environmentally stochastic IPM | | $n(z', t+1) = \int_{\alpha}^{\omega} K(z', z, \psi_t) n(z, t) dz$ | $\alpha$ = 2<br>$\omega$ = 25<br>$z$ = log(plant area) |
| | | $\psi_t = \{T_t, P_t, PAR_t, N_t, pH_t\}$ | $\psi$ = an array containing climate values |
| ***K*-kernel** | | $K(z', z, \psi_t) = P(z', z, \psi_t) + F(z', z, \psi_t)$ | |
| Sub-kernels | ***P*-subkernel** | $P(z', z, \psi_t) = s(z, \psi_t) * g(z', z, \psi_t)$ | |
| | ***F*-subkernel** | $F(z', z, \psi_t) = f_s(z) * fl_p(z) * germ_p(\psi_t) * sdl_s(z')$ | |
| Demographic functions | Survival | $\text{logit}(s(z, \psi_t)) = s_i + s_z * z + s_T * T_t + s_P * P_t + s_{PAR} * PAR_t + s_N * N_t + s_{pH} * pH_t$ | $s_i$ = -11.8<br>$s_z$ = 1.05<br>$s_T$ = 1.11<br>$s_P$ = 0.22<br>$s_{PAR}$ = -0.52<br>$s_N$ = -0.1<br>$s_{pH}$ = 0.11 |
| | Growth | $g(z', z, \psi_t) = \text{dnorm}(z', g_\mu(z, \psi_t), g_{sd})$ | $g_{sd}$ = 1.48 |
| | | $g_\mu(z, \psi_t) = g_z * z + g_T * T_t + g_P * P_t + g_{PAR} * PAR_t + g_N * N_t + g_{pH} * pH_t$ | $g_z$ = 1.02<br>$g_T$ = 0.65<br>$g_P$ = 0.02<br>$g_{PAR}$ = 0.59<br>$g_N$ = -0.04<br>$g_{pH}$ = 0.4 |
| | Reproduction | $f_s(z) = \exp(seed_i + seed_z * z)$ | $seed_i$ = -23.01<br>$seed_z$ = 1.32 |
| | | $\text{logit}(fl_p(z)) = fl_i + fl_z * z$ | $fl_i$ = -33.43<br>$fl_z$ = 1.68 |

| | | | |
|---|---|---|---|
| | | $\text{logit}(germ_p(\psi_t)) = germ_i + germ_T * T_t + germ_P * P_t + germ_{PAR} * (PAR_t/0.018) + germ_{pH} * pH_t$ | $germ_i$ = -11.8<br>$germ_T$ = 0.51<br>$germ_P$ = -0.02<br>$germ_{PAR}$ = -0.02<br>$germ_{pH}$ = 0.26 |
| | | $sdl_s(z') = \text{dnorm}(z', sdl_\mu, sdl_{sd})$ | $sdl_\mu$ = 10.23<br>$sdl_{sd}$ = 1.581 |
| Environment values | Mean temperature in warmest month | $T \sim \text{N}(0, 1.5)$ | |
| | Mean May precipitation | $P \sim \text{N}(0, 1.5)$ | |
| | PAR | $PAR \sim \text{N}(0, 1.5)$ | |
| | Soil Nitrogen | $N \sim \text{N}(0, 1.5)$ | |
| | Soil pH | $pH \sim \text{N}(0, 1.5)$ | |

**Supplementary Table 2.** Formulas, regressions and parameters used to construct the IPMs for *Calathea crotalifera*.

| Construction | | Model | Parameter |
|---|---|---|---|
| Density-independent environmentally stochastic IPM | | $n(z', t+1) = \int_{\alpha}^{\omega} K(z', z, \psi_t) n(z, t) dz$ | $\alpha$ = 0.57<br>$\omega$ = 11.9<br>$z$ = leaf area |
| | | $\psi_t = \{j_t, A_t\}$ | $\psi$ = an array containing climate values |
| ***K*-kernel** | | $K(z', z, \psi_t) = P(z', z, j_t, A_t) + F(z', z, j_t)$ | |
| Sub-kernels | ***P*-kernel** | $P(z', z, j_t, A_t) = s(z, j_t) * g(z', z, j_t, A_t)$ | |
| | ***F*-kernel** | $F(z', z, j_t) = r_p(z, j_t) * r_o(z, j_t) * n_f * n_s * s_s(j_t) * sdl_s(j_t) * sdl_{size}(z', j_t)$ | $n_f$ = 23<br>$n_s$ = 3 |
| Demographic functions | Survival | $\text{logit}(s(z, \psi_t)) = s_i + s_z * z + s_j * j_t + s_{z*j} * z * j_t$ | $s_i$ = -2.74<br>$s_z$ = 0.95<br>$s_j$ = 0.07<br>$s_{z*j}$ = -0.02 |
| | Growth | $g(z', z, j_t, A_t) = \text{dnorm}(z', g_\mu(z, j_t, A_t), g_{sd})$ | $g_{sd}$ = 1.53 |
| | | $g_\mu(z, j_t, A_t) = g_i + g_z * z + g_j * j_t + g_A * A_t + g_{z*j} * z * j_t + g_{z*A} * z * A_t + g_{j*A} * j_t * A_t + g_{z*j*A} * z * j_t * A_t$ | $g_i$ = 0.76<br>$g_z$ = 0.9<br>$g_j$ = 0.03<br>$g_A$ = 0.006<br>$g_{z*j}$ = -0.001<br>$g_{z*A}$ = 0.00045<br>$g_{j*A}$ = -0.0052<br>$g_{z*j*A}$ = 0.00035 |
| | Reproduction | $\text{logit}(r_p(z, j_t)) = r_{p,i} + r_{p,z} * z + r_{p,j} * j_t + r_{p,z*j} * z * j_t$ | $r_{p,i}$ = -13.23<br>$r_{p,z}$ = 1.401<br>$r_{p,j}$ = -0.213<br>$r_{p,z*j}$ = 0.043 |
| | | $r_o(z, j_t) = \exp(r_{o,i} + r_{o,z} * z + r_{o,j} * j_t + r_{o,z*j} * z * j_t)$ | $r_{o,i}$ = -6.673<br>$r_{o,z}$ = 0.829 |

| | | | |
|---|---|---|---|
| | | | $r_{o,j}$ = 0.067<br>$r_{o,z*j}$ = -0.007 |
| | | $s_s(j_t < 6) = 0.29$<br>$s_s(j_t \geq 6) = 0.32$ | |
| | | $sdl_s(j_t < 6) = 0.14$<br>$sdl_s(j_t \geq 6) = 0.95$ | |
| | | $sdl_{size}(z', j_t < 6) = \text{dnorm}(z', 3.08, 0.54)$<br>$sdl_{size}(z', j_t \geq 6) = \text{dnorm}(z', 2.88, 1.4)$ | |
| Environment values | Canopy openness* | $j \sim \text{N}(3, 1.4)$ | |
| | Photosynthetic rate* | $A \sim \text{N}(6, 0.8)$ | |

* In Westerband and Horvitz (2016), canopy openness ($j$) and photosynthetic rate ($A$) were modelled as random samples from a sequence of values or draws from a uniform distribution. Specifically canopy openness was realized at time $t$ as random draws from the sequence $\{1, 2, 3, 4, 5\}$ whilst photosynthetic rate was realized at time $t$ as random draws from a uniform distruction (*i.e.*, $A \sim \text{U}(5, 7)$). However, since our manipulation of the environment involves explicitly changing the temporal variance of a series, we coerced the distributions into normal distributions with the same mean and reported variance of the original sampling distributions reported in Westerband and Horvitz (2016).

**Supplementary Table 3.** Formulas, regressions and parameters used to construct the IPMs for *Heliconia tortuosa*.

| Construction | | Model | Parameter |
|---|---|---|---|
| Density-independent environmentally stochastic IPM | | $n(z', t+1) = \int_{\alpha}^{\omega} K(z', z, \psi_t) n(z, t) dz$ | $\alpha$ = 0.78<br>$\omega$ = 11.07<br>$z$ = leaf area |
| | | $\psi_t = \{j_t, A_t\}$ | $\psi$ = an array containing climate values |
| ***K***-kernel | | $K(z', z, \psi_t) = P(z', z, j_t, A_t) + F(z', z, j_t)$ | |
| Sub-kernels | ***P***-kernel | $P(z', z, j_t, A_t) = s(z, j_t) * g(z', z, j_t, A_t)$ | |
| | ***F***-kernel | $F(z', z, j_t) = r_p(z, j_t) * r_o(z, j_t) * n_f * n_s * s_s(j_t) * sdl_s(j_t) * sdl_{size}(z', j_t)$ | $n_f$ = 37<br>$n_s$ = 2.5 |
| Demographic functions | Survival | $\text{logit}(s(z, \psi_t)) = s_i + s_z * z + s_j * j_t + s_{z*j} * z * j_t$ | $s_i$ = -2.05<br>$s_z$ = 0.78<br>$s_j$ = -0.22<br>$s_{z*j}$ = 0.05 |
| | Growth | $g(z', z, j_t, A_t) = \text{dnorm}(z', g_\mu(z, j_t, A_t), g_{sd})$ | $g_{sd}$ = 0.71 |
| | | $g_\mu(z, j_t, A_t) = g_i + g_z * z + g_j * j_t + g_A * A_t + g_{z*j} * z * j_t + g_{z*A} * z * A_t + g_{j*A} * j_t * A_t + g_{z*j*A} * z * j_t * A_t$ | $g_i$ = 2.6<br>$g_z$ = 0.56<br>$g_j$ = -1.55<br>$g_A$ = 0.44<br>$g_{z*j}$ = 0.18<br>$g_{z*A}$ = -0.034<br>$g_{j*A}$ = 0.014<br>$g_{z*j*A}$ = -0.0014 |
| | Reproduction | $\text{logit}(r_p(z, j_t)) = r_{p,i} + r_{p,z} * z + r_{p,j} * j_t + r_{p,z*j} * z * j_t$ | $r_{p,i}$ = -12.55<br>$r_{p,z}$ = 1.527<br>$r_{p,j}$ = 0.154<br>$r_{p,z*j}$ = -0.013 |
| | | $r_o(z, j_t) = \exp(r_{o,i} + r_{o,z} * z + r_{o,j} * j_t + r_{o,z*j} * z * j_t)$ | $r_{o,i}$ = -1.009<br>$r_{o,z}$ = 0.157 |

| | | | |
|---|---|---|---|
| | | | $r_{o,j}$ = -0.382<br>$r_{o,z*j}$ = 0.048 |
| | | $s_s(j_t < 6) = 0.15$<br>$s_s(j_t \geq 6) = 0.2$ | |
| | | $sdl_s(j_t < 6) = 0.26$<br>$sdl_s(j_t \geq 6) = 0.33$ | |
| | | $sdl_{size}(z', j_t < 6) = \text{dnorm}(z', 2.73, 0.71)$<br>$sdl_{size}(z', j_t \geq 6) = \text{dnorm}(z', 2.34, 1.17)$ | |
| Environment values | Canopy openness | $j \sim \text{N}(3, 1.4)$ | |
| | Photosynthetic rate | $A \sim \text{N}(6.5, 0.8654937)$ | |

* In Westerband and Horvitz (2016), canopy openness ($j$) and photosynthetic rate ($A$) were modelled as random samples from a sequence of values or draws from a uniform distribution. Specifically canopy openness was realized at time $t$ as random draws from the sequence $\{1, 2, 3, 4, 5\}$ whilst photosynthetic rate was realized at time $t$ as random draws from a uniform distruction (*i.e.*, $A \sim \text{U}(5, 8)$). However, since our manipulation of the environment involves explicitly changing the temporal variance of a series, we coerced the distributions into normal distributions with the same mean and reported variance of the original sampling distributions reported in Westerband and Horvitz (2016).

**Standardised analysis pipeline.**

To investigate the roles of environmental autocorrelation and variance on demographic buffering, we used a standardised analysis pipeline to reproducibly model the linear and non-linear impacts of environment components (*i.e.*, environment autocorrelation and variance).

The standardised analysis pipeline has two phases. First, a suite of pre-defined statistical models were used to model the specified response variable (*e.g.*, **P**-**F** contribution). These models were classified as *a priori* (see below).

*A priori* models:

1. $response\ variable \sim \sigma^2 + \varphi + \sigma^2 * \varphi$
2. $response\ variable \sim \sigma^2 + \varphi + \sigma^2 * \varphi + \varphi^2$
3. $response\ variable \sim \sigma^2 + \varphi + \sigma^2 * \varphi + \varphi^2 + \varphi^3$
4. $response\ variable \sim \sigma^2 + \varphi + \sigma^2 * \varphi + (\sigma^2)^2$
5. $response\ variable \sim \sigma^2 + \varphi + \sigma^2 * \varphi + (\sigma^2)^2 + \varphi^2$
6. $response\ variable \sim \sigma^2 + \varphi + \sigma^2 * \varphi + (\sigma^2)^2 + \varphi^2 + \varphi^3$
7. $response\ variable \sim \sigma^2 + \varphi + \sigma^2 * \varphi + (\sigma^2)^2 + (\sigma^2)^3$
8. $response\ variable \sim \sigma^2 + \varphi + \sigma^2 * \varphi + (\sigma^2)^2 + (\sigma^2)^3 + \varphi^2$
9. $response\ variable \sim \sigma^2 + \varphi + \sigma^2 * \varphi + (\sigma^2)^2 + (\sigma^2)^3 + \varphi^2 + \varphi^3$

We chose this list of statistical models as they (1) contain linear and linear-interaction terms of environment autocorrelation ($\varphi$) and environment variance ($\sigma^2$) (see the first three terms of each model), (2) they sequentially add combinations of environment autocorrelation and variance up to a cubic term and (3) simplifies the combination of possible models generate from 7 predictors (127 possible combinations).

After identifying the statistical model with the lowest AIC out of all of the *a priori* models, the model with the lowest AIC was passed through an ANOVA to estimate the significance of each of the predictors (α=0.05). If all of the predictors were significant ($p$<0.05), the *a priori* model with the lowest AIC was chosen. However, if one or more of the predictors were not significant, further statistical models were constructed in the second phase of the pipeline.

In the second phase, all possible versions of the model selected in the first phase were constructed with the insignificant predictors knocked. For example, if *a priori* model 5 had the lowest AIC, but ($\sigma^2 * \varphi$) and ($\varphi^2$) were insignificant, the following 3 *post-hoc* models would be constructed:

10. $response\ variable \sim \sigma^2 + \varphi + (\sigma^2)^2 + \varphi^2$ [knock out $\sigma^2 * \varphi$]
11. $response\ variable \sim \sigma^2 + \varphi + \sigma^2 * \varphi + (\sigma^2)^2$ [knock out $\varphi^2$]
12. $response\ variable \sim \sigma^2 + \varphi + (\sigma^2)^2$ [knock out $\sigma^2 * \varphi$ and $\varphi^2$]

After constructing the *post-hoc* models, AIC values were calculated and the model with the lowest AIC was selected.

**Supplementary Table 4.** Model selection to quantify the effects of environmental autocorrelation and variance on demographic buffering (DB) in *Berberis thunbergii*. This model selection corresponds to the data show in Figure 2A in the main text. Environmental autocorrelation is denoted as $\varphi$ whilst environmental variance is denoted as $\sigma^2$.

| Model type | Model number | Model | DF | AIC | Initial selection | Final selection |
|---|---|---|---|---|---|---|
| *A priori* | 1 | $DB \sim \sigma^2 + \varphi + \sigma^2 * \varphi$ | 5 | -1333.377 | | |
| | 2 | $DB \sim \sigma^2 + \varphi + \sigma^2 * \varphi + \varphi^2$ | 6 | -1566.159 | | |
| | 3 | $DB \sim \sigma^2 + \varphi + \sigma^2 * \varphi + \varphi^2 + \varphi^3$ | 7 | -1681.584 | | |
| | 4 | $DB \sim \sigma^2 + \varphi + \sigma^2 * \varphi + (\sigma^2)^2$ | 6 | -1349.455 | | |
| | 5 | $DB \sim \sigma^2 + \varphi + \sigma^2 * \varphi + (\sigma^2)^2 + \varphi^2$ | 7 | -1619.828 | | |
| | 6 | $DB \sim \sigma^2 + \varphi + \sigma^2 * \varphi + (\sigma^2)^2 + \varphi^2 + \varphi^3$ | 8 | -1783.319 | **SELECTED** | |
| | 7 | $DB \sim \sigma^2 + \varphi + \sigma^2 * \varphi + (\sigma^2)^2 + (\sigma^2)^3$ | 7 | -1347.510 | | |
| | 8 | $DB \sim \sigma^2 + \varphi + \sigma^2 * \varphi + (\sigma^2)^2 + (\sigma^2)^3 + \varphi^2$ | 8 | -1618.012 | | |
| | 9 | $DB \sim \sigma^2 + \varphi + \sigma^2 * \varphi + (\sigma^2)^2 + (\sigma^2)^3 + \varphi^2 + \varphi^3$ | 9 | -1781.704 | | |
| *Post hoc* | 10 | $DB \sim \sigma^2 + \sigma^2 * \varphi + (\sigma^2)^2 + \varphi^2 + \varphi^3$ | 7 | -1785.317 | | **SELECTED** |
| Complete formula | | $DB \sim -0.48626 + 1.242738 * [\sigma^2] + 0.026047 * [\sigma^2 * \varphi] - 1.003697 * [(\sigma^2)^2] - 0.045427 * [\varphi^2] - 0.050135 * [\varphi^3]$ | | | | |

**Supplementary Table 5.** Model selection to quantify the effects of environmental autocorrelation and variance on demographic buffering (DB) in *Calathea crotalifera*. This model selection corresponds to the data show in Figure 2B in the main text. Environmental autocorrelation is denoted as $\varphi$ whilst environmental variance is denoted as $\sigma^2$.

| Model type | Model number | Model | DF | AIC | Initial selection | Final selection |
|---|---|---|---|---|---|---|
| *A priori* | 1 | $DB \sim \sigma^2 + \varphi + \sigma^2 * \varphi$ | 5 | -925.2545 | | |
| | 2 | $DB \sim \sigma^2 + \varphi + \sigma^2 * \varphi + \varphi^2$ | 6 | -953.4019 | | |
| | 3 | $DB \sim \sigma^2 + \varphi + \sigma^2 * \varphi + \varphi^2 + \varphi^3$ | 7 | -980.3957 | | |
| | 4 | $DB \sim \sigma^2 + \varphi + \sigma^2 * \varphi + (\sigma^2)^2$ | 6 | -927.4461 | | |
| | 5 | $DB \sim \sigma^2 + \varphi + \sigma^2 * \varphi + (\sigma^2)^2 + \varphi^2$ | 7 | -956.2009 | | |
| | 6 | $DB \sim \sigma^2 + \varphi + \sigma^2 * \varphi + (\sigma^2)^2 + \varphi^2 + \varphi^3$ | 8 | -983.8628 | **SELECTED** | |
| | 7 | $DB \sim \sigma^2 + \varphi + \sigma^2 * \varphi + (\sigma^2)^2 + (\sigma^2)^3$ | 7 | -925.4511 | | |
| | 8 | $DB \sim \sigma^2 + \varphi + \sigma^2 * \varphi + (\sigma^2)^2 + (\sigma^2)^3 + \varphi^2$ | 8 | -954.2067 | | |
| | 9 | $DB \sim \sigma^2 + \varphi + \sigma^2 * \varphi + (\sigma^2)^2 + (\sigma^2)^3 + \varphi^2 + \varphi^3$ | 9 | -981.8694 | | |
| *Post hoc* | 10 | $DB \sim \sigma^2 + \sigma^2 * \varphi + (\sigma^2)^2 + \varphi^2 + \varphi^3$ | 7 | -985.6362 | | |
| | 11 | $DB \sim \varphi + \sigma^2 * \varphi + (\sigma^2)^2 + \varphi^2 + \varphi^3$ | 7 | -984.4633 | | |
| | 12 | $DB \sim \sigma^2 + \varphi + (\sigma^2)^2 + \varphi^2 + \varphi^3$ | 7 | -984.7991 | | |
| | 13 | $DB \sim \sigma^2 * \varphi + (\sigma^2)^2 + \varphi^2 + \varphi^3$ | 6 | -986.2380 | | **SELECTED** |
| | 14 | $DB \sim \sigma^2 + (\sigma^2)^2 + \varphi^2 + \varphi^3$ | 6 | -974.8492 | | |

| | | | | |
|---|---|---|---|---|
| | 15 | $DB \sim \varphi + (\sigma^2)^2 + \varphi^2 + \varphi^3$ | 6 | -985.4061 |
| | 16 | $DB \sim (\sigma^2)^2 + \varphi^2 + \varphi^3$ | 5 | -975.5280 |
| Complete formula | | $DB \sim 0.347261 + 0.032254 * [\sigma^2 * \varphi] - 0.603492 * [(\sigma^2)^2] + 0.049503 * [\varphi^2] - 0.108026 * [\varphi^3]$ | | |

**Supplementary Table 6.** Model selection to quantify the effects of environmental autocorrelation and variance on demographic buffering (DB) in *Heliconia tortuosa*. This model selection corresponds to the data show in Figure 2C in the main text. Environmental autocorrelation is denoted as $\varphi$ whilst environmental variance is denoted as $\sigma^2$.

| Model type | Model number | Model | DF | AIC | Initial selection | Final selection |
|---|---|---|---|---|---|---|
| *A priori* | 1 | $DB \sim \sigma^2 + \varphi + \sigma^2 * \varphi$ | 5 | -1891.832 | | |
| | 2 | $DB \sim \sigma^2 + \varphi + \sigma^2 * \varphi + \varphi^2$ | 6 | -1913.937 | | |
| | 3 | $DB \sim \sigma^2 + \varphi + \sigma^2 * \varphi + \varphi^2 + \varphi^3$ | 7 | -1930.921 | | |
| | 4 | $DB \sim \sigma^2 + \varphi + \sigma^2 * \varphi + (\sigma^2)^2$ | 6 | -1895.561 | | |
| | 5 | $DB \sim \sigma^2 + \varphi + \sigma^2 * \varphi + (\sigma^2)^2 + \varphi^2$ | 7 | -1918.323 | | |
| | 6 | $DB \sim \sigma^2 + \varphi + \sigma^2 * \varphi + (\sigma^2)^2 + \varphi^2 + \varphi^3$ | 8 | -1935.878 | **SELECTED** | |
| | 7 | $DB \sim \sigma^2 + \varphi + \sigma^2 * \varphi + (\sigma^2)^2 + (\sigma^2)^3$ | 7 | -1893.568 | | |
| | 8 | $DB \sim \sigma^2 + \varphi + \sigma^2 * \varphi + (\sigma^2)^2 + (\sigma^2)^3 + \varphi^2$ | 8 | -1916.330 | | |
| | 9 | $DB \sim \sigma^2 + \varphi + \sigma^2 * \varphi + (\sigma^2)^2 + (\sigma^2)^3 + \varphi^2 + \varphi^3$ | 9 | -1933.886 | | |
| *Post hoc* | 10 | $DB \sim \sigma^2 + \sigma^2 * \varphi + (\sigma^2)^2 + \varphi^2 + \varphi^3$ | 7 | -1937.869 | | |
| | 11 | $DB \sim \varphi + \sigma^2 * \varphi + (\sigma^2)^2 + \varphi^2 + \varphi^3$ | 7 | -1935.573 | | |
| | 12 | $DB \sim \sigma^2 + \varphi + (\sigma^2)^2 + \varphi^2 + \varphi^3$ | 7 | -1937.878 | | |
| | 13 | $DB \sim \sigma^2 * \varphi + (\sigma^2)^2 + \varphi^2 + \varphi^3$ | 6 | -1937.564 | | |
| | 14 | $DB \sim \sigma^2 + (\sigma^2)^2 + \varphi^2 + \varphi^3$ | 6 | -1939.586 | | **SELECTED** |

| | | | | |
|---|---|---|---|---|
| | 15 | $DB \sim \varphi + (\sigma^2)^2 + \varphi^2 + \varphi^3$ | 6 | -1937.573 |
| | 16 | $DB \sim (\sigma^2)^2 + \varphi^2 + \varphi^3$ | 5 | -1939.284 |
| Complete formula | | $DB \sim -0.0534906 + 0.1897105 * [\sigma^2] - 0.1655802 * [(\sigma^2)^2] + 0.0052008 \ * [\varphi^2] - 0.0091953 \ * [\varphi^3]$ | | |

**Supplementary Table 7.** Model selection to quantify the effects of environmental autocorrelation and variance on the residuals of $\sum E_{a_{ij}}^{\sigma^2}$ and $\sum E_{a_{ij}}^{\sigma^2}|$ASD in *Berberis thunbergii*. This model selection corresponds to the data show in Figure 3B in the main text. Environmental autocorrelation is denoted as $\varphi$ whilst environmental variance is denoted as $\sigma^2$.

| Model type | Model number | Model | DF | AIC | Initial selection | Final selection |
|---|---|---|---|---|---|---|
| *A priori* | 1 | $residuals{\sim}\sigma^2 + \varphi + \sigma^2 * \varphi$ | 5 | -741.7773 | | |
| | 2 | $residuals{\sim}\sigma^2 + \varphi + \sigma^2 * \varphi + \varphi^2$ | 6 | -769.6791 | | |
| | 3 | $residuals{\sim}\sigma^2 + \varphi + \sigma^2 * \varphi + \varphi^2 + \varphi^3$ | 7 | -890.4373 | | |
| | 4 | $residuals{\sim}\sigma^2 + \varphi + \sigma^2 * \varphi + (\sigma^2)^2$ | 6 | -744.6596 | | |
| | 5 | $residuals{\sim}\sigma^2 + \varphi + \sigma^2 * \varphi + (\sigma^2)^2 + \varphi^2$ | 7 | -773.2640 | | |
| | 6 | $residuals{\sim}\sigma^2 + \varphi + \sigma^2 * \varphi + (\sigma^2)^2 + \varphi^2 + \varphi^3$ | 8 | -898.1634 | **SELECTED** | **SELECTED*** |
| | 7 | $residuals{\sim}\sigma^2 + \varphi + \sigma^2 * \varphi + (\sigma^2)^2 + (\sigma^2)^3$ | 7 | -742.7117 | | |
| | 8 | $residuals{\sim}\sigma^2 + \varphi + \sigma^2 * \varphi + (\sigma^2)^2 + (\sigma^2)^3 + \varphi^2$ | 8 | -771.3237 | | |
| | 9 | $residuals{\sim}\sigma^2 + \varphi + \sigma^2 * \varphi + (\sigma^2)^2 + (\sigma^2)^3 + \varphi^2 + \varphi^3$ | 9 | -896.2684 | | |
| Complete formula | | $residuals{\sim} - 1.969071 + 3.965287 * [\sigma^2] - 0.512308 * [\varphi] + 0.364988 * [\sigma^2 * \varphi] - 1.970633 * [(\sigma^2)^2] - 0.074138 * [\varphi^2] + 0.299437 * [\varphi^3]$ | | | | |

*Since all parameters were deemed significant ($\alpha < 0.05$), no *post hoc* selection was performed.

**Supplementary Table 8.** Model selection to quantify the effects of environmental autocorrelation and variance on the residuals of $\sum E_{a_{ij}}^{\sigma^2}$ and $\sum E_{a_{ij}}^{\sigma^2}|$ASD in *Calathea crotalifera*. This model selection corresponds to the data show in Figure 3E in the main text. Environmental autocorrelation is denoted as $\varphi$ whilst environmental variance is denoted as $\sigma^2$.

| Model type | Model number | Model | DF | AIC | Initial selection | Final selection |
|---|---|---|---|---|---|---|
| *A priori* | 1 | $residuals \sim \sigma^2 + \varphi + \sigma^2 * \varphi$ | 5 | -1718.857 | | |
| | 2 | $residuals \sim \sigma^2 + \varphi + \sigma^2 * \varphi + \varphi^2$ | 6 | -1756.217 | | |
| | 3 | $residuals \sim \sigma^2 + \varphi + \sigma^2 * \varphi + \varphi^2 + \varphi^3$ | 7 | -1784.586 | | |
| | 4 | $residuals \sim \sigma^2 + \varphi + \sigma^2 * \varphi + (\sigma^2)^2$ | 6 | -1759.530 | | |
| | 5 | $residuals \sim \sigma^2 + \varphi + \sigma^2 * \varphi + (\sigma^2)^2 + \varphi^2$ | 7 | -1806.057 | | |
| | 6 | $residuals \sim \sigma^2 + \varphi + \sigma^2 * \varphi + (\sigma^2)^2 + \varphi^2 + \varphi^3$ | 8 | -1843.012 | **SELECTED** | **SELECTED*** |
| | 7 | $residuals \sim \sigma^2 + \varphi + \sigma^2 * \varphi + (\sigma^2)^2 + (\sigma^2)^3$ | 7 | -1757.962 | | |
| | 8 | $residuals \sim \sigma^2 + \varphi + \sigma^2 * \varphi + (\sigma^2)^2 + (\sigma^2)^3 + \varphi^2$ | 8 | -1804.593 | | |
| | 9 | $residuals \sim \sigma^2 + \varphi + \sigma^2 * \varphi + (\sigma^2)^2 + (\sigma^2)^3 + \varphi^2 + \varphi^3$ | 9 | -1841.650 | | |
| Complete formula | | $residuals \sim 0.638800 - 1.276380 * [\sigma^2] - 0.019216 * [\varphi] + 0.029041 * [\sigma^2 * \varphi] + 0.637492 * [(\sigma^2)^2] - 0.009601 * [\varphi^2] - 0.018321 * [\varphi^3]$ | | | | |

*Since all parameters were deemed significant ($\alpha < 0.05$), no *post hoc* selection was performed.

**Supplementary Table 9.** Model selection to quantify the effects of environmental autocorrelation and variance on the residuals of $\sum E_{a_{ij}}^{\sigma^2}$ and $\sum E_{a_{ij}}^{\sigma^2}$|ASD in *Heliconia tortuosa*. This model selection corresponds to the data show in Figure 3H in the main text. Environmental autocorrelation is denoted as $\varphi$ whilst environmental variance is denoted as $\sigma^2$.

| Model type | Model number | Model | DF | AIC | Initial selection | Final selection |
|---|---|---|---|---|---|---|
| *A priori* | 1 | $residuals \sim \sigma^2 + \varphi + \sigma^2 * \varphi$ | 5 | -1380.828 | | |
| | 2 | $residuals \sim \sigma^2 + \varphi + \sigma^2 * \varphi + \varphi^2$ | 6 | -1386.508 | | |
| | 3 | $residuals \sim \sigma^2 + \varphi + \sigma^2 * \varphi + \varphi^2 + \varphi^3$ | 7 | -1473.794 | | |
| | 4 | $residuals \sim \sigma^2 + \varphi + \sigma^2 * \varphi + (\sigma^2)^2$ | 6 | -1392.269 | | |
| | 5 | $residuals \sim \sigma^2 + \varphi + \sigma^2 * \varphi + (\sigma^2)^2 + \varphi^2$ | 7 | -1398.431 | | |
| | 6 | $residuals \sim \sigma^2 + \varphi + \sigma^2 * \varphi + (\sigma^2)^2 + \varphi^2 + \varphi^3$ | 8 | -1492.824 | **SELECTED** | |
| | 7 | $residuals \sim \sigma^2 + \varphi + \sigma^2 * \varphi + (\sigma^2)^2 + (\sigma^2)^3$ | 7 | -1391.522 | | |
| | 8 | $residuals \sim \sigma^2 + \varphi + \sigma^2 * \varphi + (\sigma^2)^2 + (\sigma^2)^3 + \varphi^2$ | 8 | -1397.730 | | |
| | 9 | $residuals \sim \sigma^2 + \varphi + \sigma^2 * \varphi + (\sigma^2)^2 + (\sigma^2)^3 + \varphi^2 + \varphi^3$ | 9 | -1492.822 | | |
| *Post hoc* | 10 | $residuals \sim \sigma^2 + \sigma^2 * \varphi + (\sigma^2)^2 + \varphi^2 + \varphi^3$ | 7 | -1493.981 | | **SELECTED** |
| Complete formula | | $residuals \sim -0.804339 + 1.592423 * [\sigma^2] - 0.065901 * [\sigma^2 * \varphi] - 0.782833 * [(\sigma^2)^2] - 0.009305 * [\varphi^2] + 0.066305 * [\varphi^3]$ | | | | |

**Supplementary Table 10.** Model selection to quantify the effects of environmental autocorrelation and variance on the mean buffered size in *Berberis thunbergii*. This model selection corresponds to the data show in Figure 3C in the main text. Environmental autocorrelation is denoted as $\varphi$ whilst environmental variance is denoted as $\sigma^2$.

| Model type | Model number | Model | DF | AIC | Initial selection | Final selection |
|---|---|---|---|---|---|---|
| *A priori* | 1 | $mean\ buffered\ size{\sim}\sigma^2 + \varphi + \sigma^2 * \varphi$ | 5 | -1808.928 | | |
| | 2 | $mean\ buffered\ size{\sim}\sigma^2 + \varphi + \sigma^2 * \varphi + \varphi^2$ | 6 | -2170.820 | | |
| | 3 | $mean\ buffered\ size{\sim}\sigma^2 + \varphi + \sigma^2 * \varphi + \varphi^2 + \varphi^3$ | 7 | -2291.923 | **SELECTED** | **SELECTED** |
| | 4 | $mean\ buffered\ size{\sim}\sigma^2 + \varphi + \sigma^2 * \varphi + (\sigma^2)^2$ | 6 | -1807.061 | | |
| | 5 | $mean\ buffered\ size{\sim}\sigma^2 + \varphi + \sigma^2 * \varphi + (\sigma^2)^2 + \varphi^2$ | 7 | -2169.493 | | |
| | 6 | $mean\ buffered\ size{\sim}\sigma^2 + \varphi + \sigma^2 * \varphi + (\sigma^2)^2 + \varphi^2 + \varphi^3$ | 8 | -2291.086 | | |
| | 7 | $mean\ buffered\ size{\sim}\sigma^2 + \varphi + \sigma^2 * \varphi + (\sigma^2)^2 + (\sigma^2)^3$ | 7 | -1805.061 | | |
| | 8 | $mean\ buffered\ size{\sim}\sigma^2 + \varphi + \sigma^2 * \varphi + (\sigma^2)^2 + (\sigma^2)^3 + \varphi^2$ | 8 | -2167.493 | | |
| | 9 | $mean\ buffered\ size{\sim}\sigma^2 + \varphi + \sigma^2 * \varphi + (\sigma^2)^2 + (\sigma^2)^3 + \varphi^2 + \varphi^3$ | 9 | -2289.087 | | |
| *Post hoc* | 10 | $mean\ buffered\ size{\sim}\sigma^2 + \sigma^2 * \varphi + \varphi^2 + \varphi^3$ | 8 | -2288.461 | | |
| | 11 | $mean\ buffered\ size{\sim}\sigma^2 + \varphi + \varphi^2 + \varphi^3$ | 8 | -2290.699 | | |
| | 12 | $mean\ buffered\ size{\sim}\sigma^2 + \varphi^2 + \varphi^3$ | 7 | -2144.997 | | |
| Complete formula | | $mean\ buffered\ size{\sim}0.6679336 - 0.0313268 * [\sigma^2] - 0.0051673 * [\varphi] - 0.0019560 * [\sigma^2 * \varphi] - 0.0175639 * [\varphi^2] + 0.0133206 * [\varphi^3]$ | | | | |

**Supplementary Table 11.** Model selection to quantify the effects of environmental autocorrelation and variance on the mean buffered size in *Calathea crotalifera*. This model selection corresponds to the data show in Figure 3F in the main text. Environmental autocorrelation is denoted as $\varphi$ whilst environmental variance is denoted as $\sigma^2$.

| Model type | Model number | Model | DF | AIC | Initial selection | Final selection |
|---|---|---|---|---|---|---|
| *A priori* | 1 | $mean\ buffered\ size \sim \sigma^2 + \varphi + \sigma^2 * \varphi$ | 5 | -3226.940 | | |
| | 2 | $mean\ buffered\ size \sim \sigma^2 + \varphi + \sigma^2 * \varphi + \varphi^2$ | 6 | -3241.007 | | |
| | 3 | $mean\ buffered\ size \sim \sigma^2 + \varphi + \sigma^2 * \varphi + \varphi^2 + \varphi^3$ | 7 | -3261.367 | **SELECTED** | **SELECTED** |
| | 4 | $mean\ buffered\ size \sim \sigma^2 + \varphi + \sigma^2 * \varphi + (\sigma^2)^2$ | 6 | -3225.148 | | |
| | 5 | $mean\ buffered\ size \sim \sigma^2 + \varphi + \sigma^2 * \varphi + (\sigma^2)^2 + \varphi^2$ | 7 | -3239.230 | | |
| | 6 | $mean\ buffered\ size \sim \sigma^2 + \varphi + \sigma^2 * \varphi + (\sigma^2)^2 + \varphi^2 + \varphi^3$ | 8 | -3259.614 | | |
| | 7 | $mean\ buffered\ size \sim \sigma^2 + \varphi + \sigma^2 * \varphi + (\sigma^2)^2 + (\sigma^2)^3$ | 7 | -3223.159 | | |
| | 8 | $mean\ buffered\ size \sim \sigma^2 + \varphi + \sigma^2 * \varphi + (\sigma^2)^2 + (\sigma^2)^3 + \varphi^2$ | 8 | -3237.243 | | |
| | 9 | $mean\ buffered\ size \sim \sigma^2 + \varphi + \sigma^2 * \varphi + (\sigma^2)^2 + (\sigma^2)^3 + \varphi^2 + \varphi^3$ | 9 | -3257.628 | | |
| *Post hoc* | 10 | $mean\ buffered\ size \sim \sigma^2 + \sigma^2 * \varphi + \varphi^2 + \varphi^3$ | 8 | -3252.451 | | |
| | 11 | $mean\ buffered\ size \sim \sigma^2 + \varphi + \varphi^2 + \varphi^3$ | 8 | -3249.336 | | |
| | 12 | $mean\ buffered\ size \sim \sigma^2 + \varphi^2 + \varphi^3$ | 7 | -3241.391 | | |
| Complete formula | | $mean\ buffered\ size \sim 0.8956762 + 0.0007139 * [\sigma^2] + 0.0009980 * [\varphi] - 0.0011847 * [\sigma^2 * \varphi] + 0.0002204 * [\varphi^2] + 0.0005852 * [\varphi^3]$ | | | | |

**Supplementary Table 12.** Model selection to quantify the effects of environmental autocorrelation and variance on the mean buffered size in *Heliconia tortuosa*. This model selection corresponds to the data show in Figure 3I in the main text. Environmental autocorrelation is denoted as $\varphi$ whilst environmental variance is denoted as $\sigma^2$.

| Model type | Model number | Model | DF | AIC | Initial selection | Final selection |
|---|---|---|---|---|---|---|
| *A priori* | 1 | $mean\ buffered\ size \sim \sigma^2 + \varphi + \sigma^2 * \varphi$ | 5 | -1971.458 | | |
| | 2 | $mean\ buffered\ size \sim \sigma^2 + \varphi + \sigma^2 * \varphi + \varphi^2$ | 6 | -2090.884 | | |
| | 3 | $mean\ buffered\ size \sim \sigma^2 + \varphi + \sigma^2 * \varphi + \varphi^2 + \varphi^3$ | 7 | -2119.626 | **SELECTED** | **SELECTED*** |
| | 4 | $mean\ buffered\ size \sim \sigma^2 + \varphi + \sigma^2 * \varphi + (\sigma^2)^2$ | 6 | -1970.073 | | |
| | 5 | $mean\ buffered\ size \sim \sigma^2 + \varphi + \sigma^2 * \varphi + (\sigma^2)^2 + \varphi^2$ | 7 | -2089.940 | | |
| | 6 | $mean\ buffered\ size \sim \sigma^2 + \varphi + \sigma^2 * \varphi + (\sigma^2)^2 + \varphi^2 + \varphi^3$ | 8 | -2118.838 | | |
| | 7 | $mean\ buffered\ size \sim \sigma^2 + \varphi + \sigma^2 * \varphi + (\sigma^2)^2 + (\sigma^2)^3$ | 7 | -1968.075 | | |
| | 8 | $mean\ buffered\ size \sim \sigma^2 + \varphi + \sigma^2 * \varphi + (\sigma^2)^2 + (\sigma^2)^3 + \varphi^2$ | 8 | -2087.944 | | |
| | 9 | $mean\ buffered\ size \sim \sigma^2 + \varphi + \sigma^2 * \varphi + (\sigma^2)^2 + (\sigma^2)^3 + \varphi^2 + \varphi^3$ | 9 | -2116.842 | | |
| Complete formula | | $mean\ buffered\ size \sim 0.8411429 + 0.0082338 * [\sigma^2] - 0.1178307 * [\varphi] + 0.0912694 * [\sigma^2 * \varphi] + 0.0088286 * [\varphi^2] + 0.0087587 * [\varphi^3]$ | | | | |

*Since all parameters were deemed significant ($\alpha < 0.05$), no *post hoc* selection was performed.

**Supplementary Table 13.** Model selection to quantify the effects of environmental autocorrelation and variance on the **P-F** contribution in *Berberis thunbergii*. This model selection corresponds to the data show in Figure 4B in the main text. Environmental autocorrelation is denoted as $\varphi$ whilst environmental variance is denoted as $\sigma^2$.

| Model type | Model number | Model | DF | AIC | Initial selection | Final selection |
|---|---|---|---|---|---|---|
| *A priori* | 1 | $\mathbf{P-F}\ contribution \sim \sigma^2 + \varphi + \sigma^2 * \varphi$ | 5 | -8282.140 | | |
| | 2 | $\mathbf{P-F}\ contribution \sim \sigma^2 + \varphi + \sigma^2 * \varphi + \varphi^2$ | 6 | -8747.482 | | |
| | 3 | $\mathbf{P-F}\ contribution \sim \sigma^2 + \varphi + \sigma^2 * \varphi + \varphi^2 + \varphi^3$ | 7 | -8783.395 | | |
| | 4 | $\mathbf{P-F}\ contribution \sim \sigma^2 + \varphi + \sigma^2 * \varphi + (\sigma^2)^2$ | 6 | -8286.741 | | |
| | 5 | $\mathbf{P-F}\ contribution \sim \sigma^2 + \varphi + \sigma^2 * \varphi + (\sigma^2)^2 + \varphi^2$ | 7 | -8804.515 | | |
| | 6 | $\mathbf{P-F}\ contribution \sim \sigma^2 + \varphi + \sigma^2 * \varphi + (\sigma^2)^2 + \varphi^2 + \varphi^3$ | 8 | -8853.171 | **SELECTED** | |
| | 7 | $\mathbf{P-F}\ contribution \sim \sigma^2 + \varphi + \sigma^2 * \varphi + (\sigma^2)^2 + (\sigma^2)^3$ | 7 | -8284.763 | | |
| | 8 | $\mathbf{P-F}\ contribution \sim \sigma^2 + \varphi + \sigma^2 * \varphi + (\sigma^2)^2 + (\sigma^2)^3 + \varphi^2$ | 8 | -8802.731 | | |
| | 9 | $\mathbf{P-F}\ contribution \sim \sigma^2 + \varphi + \sigma^2 * \varphi + (\sigma^2)^2 + (\sigma^2)^3 + \varphi^2 + \varphi^3$ | 9 | -8851.442 | | |
| *Post hoc* | 10 | $\mathbf{P-F}\ contribution \sim \sigma^2 + \sigma^2 * \varphi + (\sigma^2)^2 + \varphi^2 + \varphi^3$ | 7 | -8853.432 | | **SELECTED** |
| Complete formula | | $\mathbf{P-F}\ contribution \sim [5.943e-08] - [1.541e-07] * [\sigma^2] - [4.113e-09] * [\sigma^2 * \varphi] + [1.208e-07] * [(\sigma^2)^2] + [1.038e-08] * [\varphi^2] + [3.724e-09] * [\varphi^3]$ | | | | |

**Supplementary Table 14.** Model selection to quantify the effects of environmental autocorrelation and variance on the **P-F** contribution in *Calathea crotalifera*. This model selection corresponds to the data show in Figure 4C in the main text. Environmental autocorrelation is denoted as $\varphi$ whilst environmental variance is denoted as $\sigma^2$.

| Model type | Model number | Model | DF | AIC | Initial selection | Final selection |
|---|---|---|---|---|---|---|
| *A priori* | 1 | $\mathbf{P-F}\ contribution \sim \sigma^2 + \varphi + \sigma^2 * \varphi$ | 5 | -8011.995 | | |
| | 2 | $\mathbf{P-F}\ contribution \sim \sigma^2 + \varphi + \sigma^2 * \varphi + \varphi^2$ | 6 | -8040.051 | | |
| | 3 | $\mathbf{P-F}\ contribution \sim \sigma^2 + \varphi + \sigma^2 * \varphi + \varphi^2 + \varphi^3$ | 7 | -8067.034 | | |
| | 4 | $\mathbf{P-F}\ contribution \sim \sigma^2 + \varphi + \sigma^2 * \varphi + (\sigma^2)^2$ | 6 | -8014.190 | | |
| | 5 | $\mathbf{P-F}\ contribution \sim \sigma^2 + \varphi + \sigma^2 * \varphi + (\sigma^2)^2 + \varphi^2$ | 7 | -8042.851 | | |
| | 6 | $\mathbf{P-F}\ contribution \sim \sigma^2 + \varphi + \sigma^2 * \varphi + (\sigma^2)^2 + \varphi^2 + \varphi^3$ | 8 | -8070.503 | **SELECTED** | |
| | 7 | $\mathbf{P-F}\ contribution \sim \sigma^2 + \varphi + \sigma^2 * \varphi + (\sigma^2)^2 + (\sigma^2)^3$ | 7 | -8012.195 | | |
| | 8 | $\mathbf{P-F}\ contribution \sim \sigma^2 + \varphi + \sigma^2 * \varphi + (\sigma^2)^2 + (\sigma^2)^3 + \varphi^2$ | 8 | -8040.857 | | |
| | 9 | $\mathbf{P-F}\ contribution \sim \sigma^2 + \varphi + \sigma^2 * \varphi + (\sigma^2)^2 + (\sigma^2)^3 + \varphi^2 + \varphi^3$ | 9 | -8068.509 | | |
| *Post hoc* | 10 | $\mathbf{P-F}\ contribution \sim \sigma^2 + \sigma^2 * \varphi + (\sigma^2)^2 + \varphi^2 + \varphi^3$ | 7 | -8072.276 | | |
| | 11 | $\mathbf{P-F}\ contribution \sim \varphi + \sigma^2 * \varphi + (\sigma^2)^2 + \varphi^2 + \varphi^3$ | 7 | -8071.103 | | |
| | 12 | $\mathbf{P-F}\ contribution \sim \sigma^2 + \varphi + (\sigma^2)^2 + \varphi^2 + \varphi^3$ | 7 | -8071.438 | | |
| | 13 | $\mathbf{P-F}\ contribution \sim \varphi + (\sigma^2)^2 + \varphi^2 + \varphi^3$ | 6 | -8072.045 | | |
| | 14 | $\mathbf{P-F}\ contribution \sim \sigma^2 + (\sigma^2)^2 + \varphi^2 + \varphi^3$ | 6 | -8061.454 | | |

| | | | | | | |
|---|---|---|---|---|---|---|
| | 15 | $\mathbf{P-F}\ contribution \sim \sigma^2 * \varphi + (\sigma^2)^2 + \varphi^2 + \varphi^3$ | 6 | -8072.878 | | **SELECTED** |
| | 16 | $\mathbf{P-F}\ contribution \sim (\sigma^2)^2 + \varphi^2 + \varphi^3$ | 5 | -8062.133 | | |
| Complete formula | | $\mathbf{P-F}\ contribution \sim [5.029e-08] - [8.739e-08] * [(\sigma^2)^2] + 4.676e-09 * [\sigma^2 * \varphi] + [7.155e-09] * [\varphi^2] - [1.564e-08] * [\varphi^3]$ | | | | |

**Supplementary Table 15.** Model selection to quantify the effects of environmental autocorrelation and variance on the **P-F** contribution in *Heliconia tortuosa*. This model selection corresponds to the data show in Figure 4D in the main text. Environmental autocorrelation is denoted as $\varphi$ whilst environmental variance is denoted as $\sigma^2$.

| Model type | Model number | Model | DF | AIC | Initial selection | Final selection |
|---|---|---|---|---|---|---|
| *A priori* | 1 | $\mathbf{P-F}\ contribution \sim \sigma^2 + \varphi + \sigma^2 * \varphi$ | 5 | -8958.788 | | |
| | 2 | $\mathbf{P-F}\ contribution \sim \sigma^2 + \varphi + \sigma^2 * \varphi + \varphi^2$ | 6 | -8986.418 | | |
| | 3 | $\mathbf{P-F}\ contribution \sim \sigma^2 + \varphi + \sigma^2 * \varphi + \varphi^2 + \varphi^3$ | 7 | -9010.538 | | |
| | 4 | $\mathbf{P-F}\ contribution \sim \sigma^2 + \varphi + \sigma^2 * \varphi + (\sigma^2)^2$ | 6 | -8962.007 | | |
| | 5 | $\mathbf{P-F}\ contribution \sim \sigma^2 + \varphi + \sigma^2 * \varphi + (\sigma^2)^2 + \varphi^2$ | 7 | -8990.381 | | |
| | 6 | $\mathbf{P-F}\ contribution \sim \sigma^2 + \varphi + \sigma^2 * \varphi + (\sigma^2)^2 + \varphi^2 + \varphi^3$ | 8 | -9015.246 | **SELECTED** | |
| | 7 | $\mathbf{P-F}\ contribution \sim \sigma^2 + \varphi + \sigma^2 * \varphi + (\sigma^2)^2 + (\sigma^2)^3$ | 7 | -8960.023 | | |
| | 8 | $\mathbf{P-F}\ contribution \sim \sigma^2 + \varphi + \sigma^2 * \varphi + (\sigma^2)^2 + (\sigma^2)^3 + \varphi^2$ | 8 | -8988.399 | | |
| | 9 | $\mathbf{P-F}\ contribution \sim \sigma^2 + \varphi + \sigma^2 * \varphi + (\sigma^2)^2 + (\sigma^2)^3 + \varphi^2 + \varphi^3$ | 9 | -9013.267 | | |
| *Post hoc* | 10 | $\mathbf{P-F}\ contribution \sim \sigma^2 + \sigma^2 * \varphi + (\sigma^2)^2 + \varphi^2 + \varphi^3$ | 7 | -9017.245 | | **SELECTED** |
| | 11 | $\mathbf{P-F}\ contribution \sim \varphi + \sigma^2 * \varphi + (\sigma^2)^2 + \varphi^2 + \varphi^3$ | 7 | -9015.157 | | |
| | 12 | $\mathbf{P-F}\ contribution \sim \sigma^2 + \varphi + (\sigma^2)^2 + \varphi^2 + \varphi^3$ | 7 | -9016.691 | | |
| | 13 | $\mathbf{P-F}\ contribution \sim \varphi + (\sigma^2)^2 + \varphi^2 + \varphi^3$ | 6 | -9016.607 | | |
| | 14 | $\mathbf{P-F}\ contribution \sim \sigma^2 + (\sigma^2)^2 + \varphi^2 + \varphi^3$ | 6 | -8999.407 | | |

| | | | | |
|---|---|---|---|---|
| | 15 | $\mathbf{P-F}\ contribution \sim \sigma^2 * \varphi + (\sigma^2)^2 + \varphi^2 + \varphi^3$ | 6 | -9017.155 |
| | 16 | $\mathbf{P-F}\ contribution \sim (\sigma^2)^2 + \varphi^2 + \varphi^3$ | 5 | -8999.493 |
| Complete formula | | $\mathbf{P-F}\ contribution \sim [5.029e-08] - [8.739e-08] * [(\sigma^2)^2] + 4.676e-09 * [\sigma^2 * \varphi] + [7.155e-09] * [\varphi^2] - [1.564e-08] * [\varphi^3]$ | | |
